# T cell circuits that sense antigen density with an ultrasensitive threshold

**DOI:** 10.1101/2021.01.21.427654

**Authors:** Rogelio A. Hernandez-Lopez, Wei Yu, Katie Cabral, Olivia Creasey, Maria del Pilar Lopez Pazmino, Yurie Tonai, Arsenia De Guzman, Anna Mäkelä, Kalle Saksela, Zev J. Gartner, Wendell A. Lim

## Abstract

Overexpressed tumor associated antigens (e.g. HER2 and EGFR) are attractive targets for therapeutic T cells, but toxic cross-reaction with normal tissues expressing low antigen levels has been observed with Chimeric Antigen Receptor (CAR) T cells targeting such antigens. Inspired by natural ultrasensitive response circuits, we engineer a two-step positive feedback circuit that allows T cells to discriminate targets based on a sigmoidal antigen density threshold. In this circuit, a low affinity SynNotch receptor for HER2 controls the expression of a high affinity CAR for HER2. Increasing HER2 density thus has cooperative effects on T cells ╌ it both increases CAR expression and activation ╌ leading to a sigmoidal response. T Cells with this circuit show sharp discrimination between target cells expressing normal and disease levels of HER2, both *in vitro* and *in vivo*.

**One Sentence Summary:** A two-step positive feedback circuit generates engineered T cells capable of killing target cells with an ultrasensitive antigen density threshold.

## MAIN TEXT

The specificity with which chimeric antigen receptor (CAR) T cells can recognize and kill tumor cells and discriminate against normal cells remains very limited (*1-3*). One of the major challenges is finding surface proteins that are absolutely tumor specific (*4*). Despite the remarkable recent success of using CAR T cells to treat hematologic cancers (*5-7*), these CAR T cells indiscriminately kill both cancerous and normal B cells, because they both express the target antigen CD19. While targeting CD19 is possible because B cells aplasia is tolerable, solid cancers have proven far more difficult to treat with CAR T cells, since indiscriminate killing of normal tissue can be lethal (*8, 9, 10*). For example, antigens such as HER2 and EGFR are often found overexpressed in several types of cancers, but they are also expressed at lower densities in normal epithelial tissues (*11, 12*). Not surprisingly, clinical studies with anti-HER2 CAR T cells have reported lethal cross-reaction with normal organs (*8*). This kind of on-target, off-tumor toxicity has been observed for CARs directed at several other overexpressed tumor associated antigens (*13-15*). Thus, an unresolved question on cancer immunotherapy is how to engineer T cells capable of robust discrimination of cancer cells from normal cells based on antigen density (**Fig. 1A** top). The ability to engineer such T cells would significantly improve treatment of several cancers including ovarian, gastric, esophageal and breast carcinomas.

**Figure 1.**
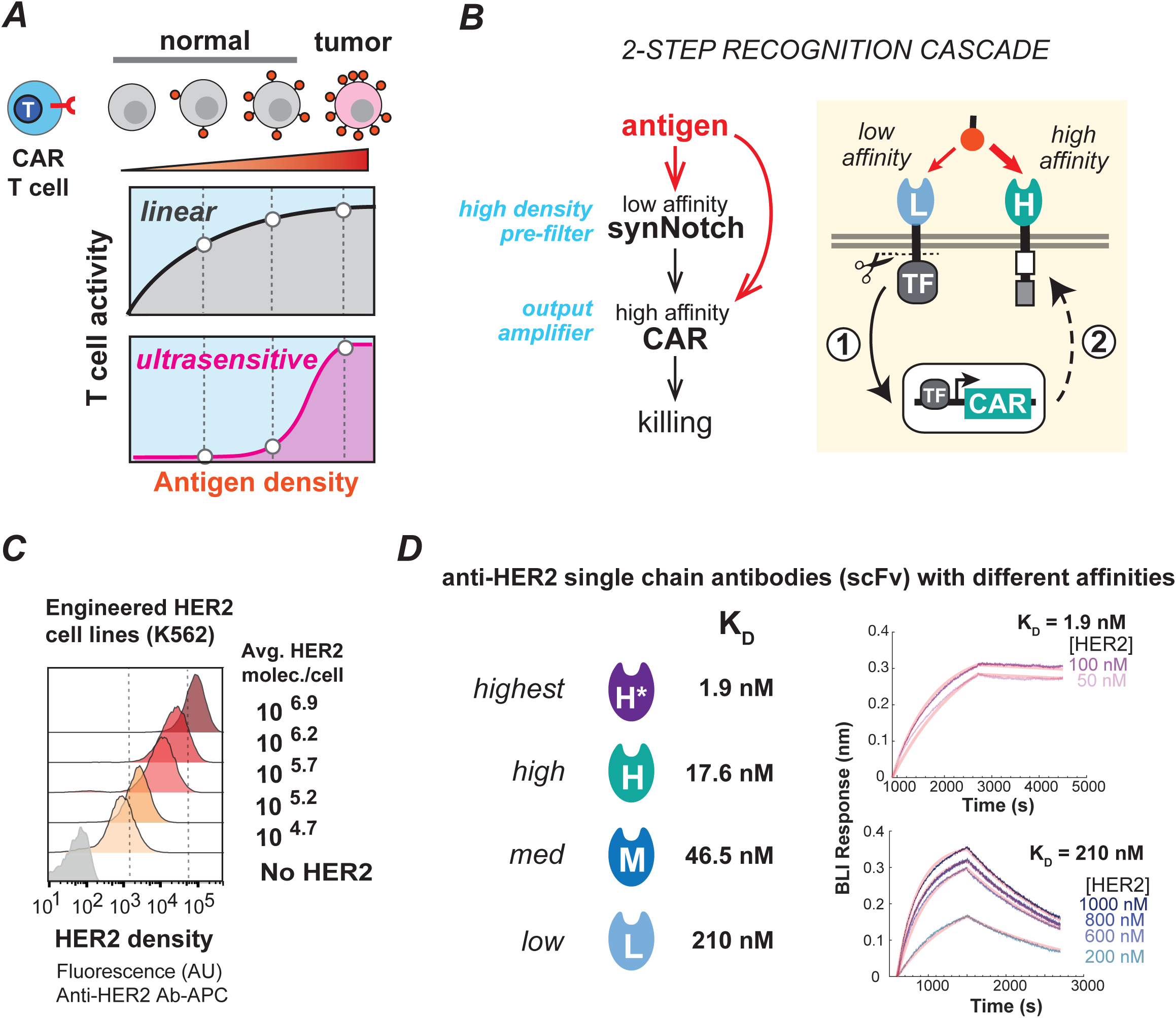
Strategy to engineer T cells capable of ultrasensitive antigen density sensing. **A**. The ideal therapeutic T cells clearly distinguishes between high antigen density expressing tumor cells and normal cells that express lower levels of a tumor associated antigen. A CAR T cell with a standard linear response curve would distinguish poorly between high- and low density cells, while good discrimination requires a sigmoidal ultrasensitive dose-response curve. **B**. Design of two-step CAR T circuit. A SynNotch receptor detects antigen (HER2) with low affinity. This SynNotch receptor, when fully activated, induces expression of a high affinity CAR. In principle, cells with this cascade combines two different elements ╌ a low affinity SynNotch that acts as a high antigen density pre-filter, and a high affinity CAR activity that would activate T cell killing and acts as an amplifier. **C**. To quantitatively assay antigen density sensing, we engineered stable cell-lines of K562 cells expressing the shown densities of the cancer associated antigen HER2. These cell lines can be compared to tumor cell lines (**Fig. S1**). The average HER2 molecules per cell was measured as shown in **Fig. S1**. To construct different HER2 sensing systems, we utilized a series of anti-HER2 single chain antibodies (scFvs) with affinities spanning a range of over 100-fold. **D**. Binding affinities for series of anti-HER2 scFvs utilized in this study (See **Fig. S2** for details about sequences and binding affinity measurements). Biolayer interferometry sensograms showing the binding kinetics for human HER2 and immobilized Anti-HER2 ScFvs. Data are shown as colored lines and the best fit for data to a 1:1 binding model is shown in light red. HER2 concentrations utilized for binding affinity measurements are indicated.

The ideal therapeutic T cell senses target antigen density and kills with an ultrasensitive response curve, characterized by a sigmoidal shape with a sharp activation threshold (**Fig. 1A** - bottom dose response plot). Sigmoidal or non-linear dose response behaviors are fundamentally important throughout biology (*16, 17*) as many critical regulatory systems require large changes in output activity in response to small changes in input level (hence the term “ultrasensitive”). This type of sigmoidal all-or-none activation curve for a CAR T cell would, in principle, yield far cleaner discrimination between cells with antigen densities below and above a desired threshold.

How might we engineer CAR T cells to recognize their antigens with a non-linear ultrasensitive density threshold? The field of systems biology has uncovered several common mechanisms for generating ultrasensitive behaviors that have been used repeatedly over the course of evolution (*18-20*). For example, cooperative binding of ligand among domains of multivalent proteins can yield molecular scale ultrasensitivity. A classic example is hemoglobin, which binds oxygen with an ultrasensitive dependence on oxygen pressure. This sigmoidal response allows the 4 monomers of hemoglobin to either fully bind or fully release oxygen based on the relatively small difference in oxygen levels in the lungs vs peripheral tissues. While it may be possible to engineer cooperatively acting CARs for T cells, this would require fairly complex and nuanced molecular engineering of key functional domains to reprogram T cell function (*21*). Efforts to engineer density sensitive CAR T cells have focused on tuning biophysical properties of CAR functional domains, such as lowering the affinity of the antibody domain or lowering the expression of the CAR construct (*22-25*). CARs with low affinity recognition domains can show moderately reduced toxicity against low antigen target cells (*22-25*). Nonetheless, such changes are predicted to shift the response curve in a linear fashion, and thus might yield limited increases in therapeutic discrimination (i.e. they also yield a lower degree of activation, leading to reduced efficacy).

An alternative commonly observed mechanism for achieving ultrasensitivity is through regulatory cascades or circuits (*18, 26, 27*). These ultrasensitive cell circuits convert linear inputs into switch-like non-linear outputs. Positive feedback cascades lead to ultrasensitive activation in many biological systems (*16*), for example certain mitogen activated protein (MAP) kinase cascades (*18, 28*) and entry to mitosis (*29*). Another example of a regulatory cascade can be found in T cells sensing the cytokine IL-2 ╌ binding of IL-2 to the basally expressed low affinity receptors results in induced expression of the high affinity alpha subunit of the IL-2 receptor (CD25) (*30*). This second transcriptional step leads to higher IL-2 sensitivity, i.e. a form of positive feedback. Here, IL-2 acts on the pathway in two ways ╌ it triggers the initiating response to express high affinity receptors, then it directly activates the high affinity receptors.

We hypothesized that we could engineer ultrasensitive positive feedback circuits for antigen density sensing with the modular synthetic biology tools that we have for immune cell engineering (*31-33*). A simple way to do this would be to first recognize the cognate antigen with a low affinity SynNotch receptor which in turn would induce expression of a high affinity CAR for the same antigen (**Fig. 1B**) (upon engaging their cognate antigen, SynNotch receptors activate transcription of a genetically encoded payload) (*32-34*). This two-step, two-receptor circuit would recognize the same antigen at two different stages in the pathway (first, antigen binding induces CAR expression; the subsequent antigen binding step activates the CAR and T cell killing) (**Fig. 1B**), and thus the response is predicted to show increased cooperativity (**Fig. 1A** bottom). In principle, changing the SynNotch and CAR affinities should also allow tunability of the antigen density response. The low affinity SynNotch receptor acts as a high antigen density pre-filter, while the high affinity CAR serves as an amplifier of robust T cell killing activity.

We chose to test this hypothesis by engineering a T cell circuit that could achieve density dependent recognition of HER2. To quantitatively analyze antigen density recognition, we constructed a series of stable K562 tumor cell lines that differ only in their level of HER2 expression over a range of ∼10^2^-fold (**Fig. 1C**). Their HER2 densities span the same range of several cancer cell lines with variable HER2 expression scores (**Fig. S1A**). Given the HER2 expression levels observed in these tumor cell lines, we set a goal of being able to robustly discriminate between cells expressing 10^6.5^ (HER2 pathology score of 3+, as defined by ASCO-CAP scoring guidelines) and 10^4.5^ molecules per cell (HER2 score of 0) (see **Fig. S1A** for quantitation of average HER2 molecules per cell). To construct the two-step circuits, we built a toolkit of anti-HER2 single chain antibodies (scFv) by introducing mutations in the scFv recognition site of the 4D5 Herceptin antibody (See **Fig. S2** for sequences information). We measured the mutant scFv affinities and determined that they span a 100-fold range - with dissociation constants between 2.0 nM and 200 nM (**Fig. 1D, Fig. S2B**).

We constructed and tested several versions of the anti-HER2 SynNotch → anti-HER2 CAR circuit in human CD8+ T cells (**Fig. S3A**). To directly assay whether this circuit design displays ultrasensitive behavior we measured the antigen density dependence of target cell killing by quantitative flow cytometry. We obtained a number of circuits that showed significantly improved antigen density ultrasensitivity (Hill coefficients (n_H_) of 1.7 − 4.4) (**Fig. 2A** bottom, **Fig. S3B, Fig. S3E**). In contrast, constitutive expression of either high or low affinity CAR show modest density discrimination (**Fig. 2A** top). When a low affinity SynNotch receptor is used to control expression of a high affinity CAR receptor (purple line, **Fig. 2A**), this yields an ultrasensitive threshold that clearly discriminates between our target densities of 10^4.5^ and 10^6.5^ (**Fig. 2B**, n_H_ = 4.4, den_50_ = 10^5.5^). Reproducibly similar ultrasensitive cytotoxic behavior was observed in engineered T cells from two different donors expressing the same circuit (n_H_ = 4.1 and n_H_ = 6.5) (**Fig. S3C**).

**Figure 2.**
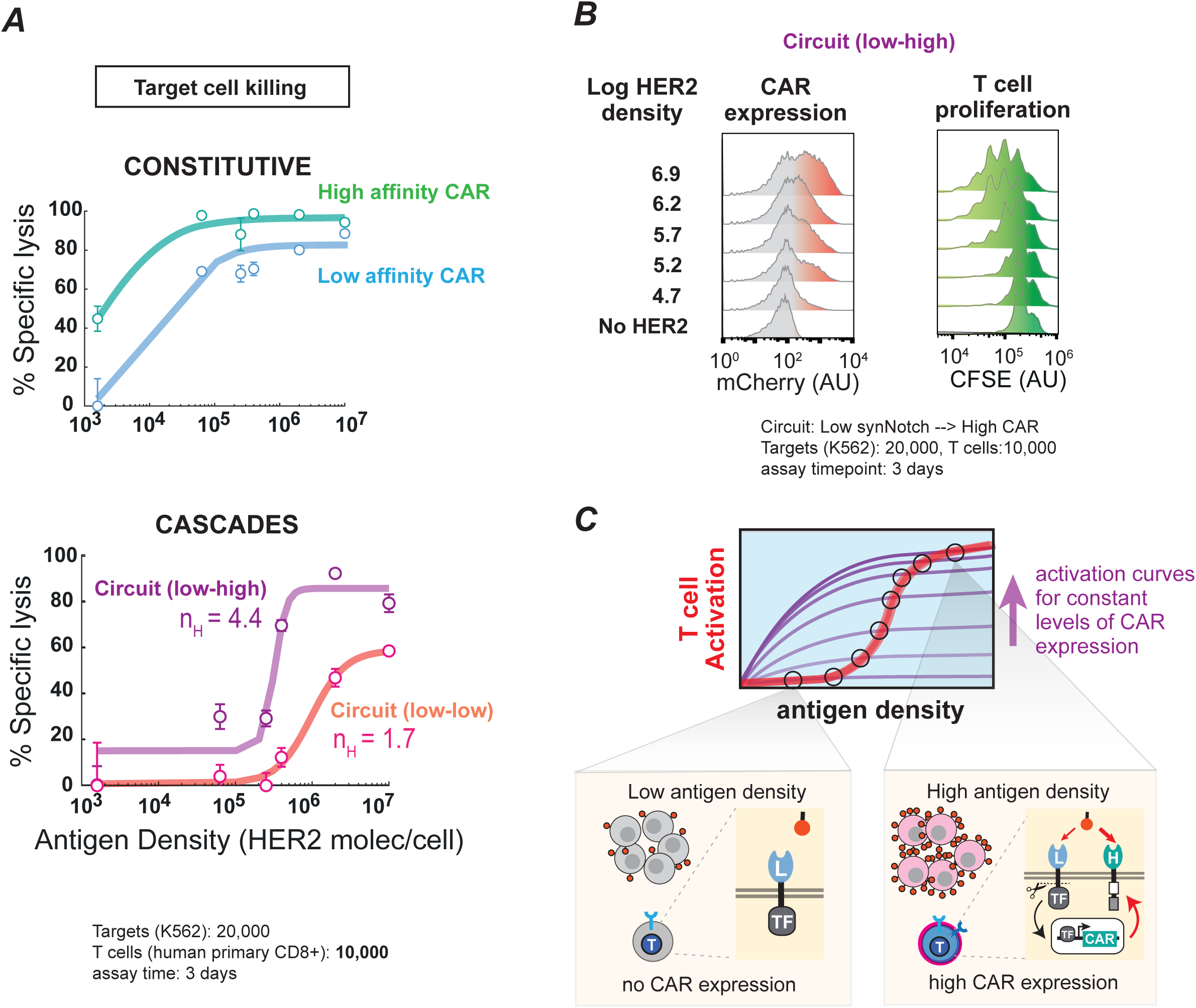
A two-step low-to-high affinity recognition circuit yields ultrasensitive antigen density sensing. **A**. In vitro cell killing curves as a function of target cell antigen density, using human primary CD8+ T cells expressing a constitutive CAR of high or low affinity (top) or a two-step circuit in which the low affinity SynNotch receptor induces expression of either a low or a high affinity CAR. For the circuits, transparent lines are fits to a hill equation (hill coefficient for each curve is indicated; see **Fig. S3B** for details). For constitutive CARs, transparent lines are drawn based on inspection. The percentage of specific lysis was determined using flow cytometry by counting the number of target cells after 3 days relative to a co-culture in the presence of untransduced T cells (see **Fig. S4A** for gating details). Data are shown as the mean and standard error from the mean (n=3). **B**. Representative FACS distributions for CAR expression and T cell proliferation measured as a function of target cell HER2 density (at 3 days) for T cells expressing a low-to-high affinity recognition circuit. As shown, significant CAR expression and T cell proliferation is only observed at HER2 densities of >10^5^. **C**. Model for the mechanism of a two-step circuit expressing a low affinity SynNotch to high affinity CAR. In principle, cells with this circuit display two very different responses ╌ in the presence of a low antigen density target (left) the T cell activity will be dominated by the low affinity SynNotch that does not express or express very low levels of a CAR. In contrast, in the presence of a high antigen density target (right) the expression of a CAR is robust, and the T cell activity is dominated by the high affinity CAR response. T cell activity is predicted to show a robust sigmoidal response curve, shown in red, because as antigen density increases, this leads to a gradual increase in CAR expression, transiting between the series of linear response curves shown in purple.

We showed that the origin of this ultrasensitivity comes from the transcriptional cascade that controls CAR expression. We measured mCherry levels (CARs were tagged with an mCherry protein, **Fig. S3A**) and observed that steady state CAR expression level depended strongly on target antigen density (**Fig. 2B** left and **Fig. S3D** red line). Moreover, induction of T cell proliferation, a critical component of an antitumor response, also shows an ultrasensitive dependence on target antigen density (**Fig. 2B** right and **Fig. S3D** green line).

The response of the T cell circuit can be tuned by altering the component affinities within the circuit. For example, lowering the CAR affinity (while maintaining a low affinity synNotch) increases the density threshold (**Fig. 2A**) (den_50_ = 10^6.0^), but also reduces the maximal killing activity and lowers the overall ultrasensitivity (n_H_ = 1.6). Ultrasensitivity, also begins to breakdown if we use a high affinity SynNotch receptor as the circuit pre-filter (**Fig. S3E**). Together, our data shows that the ultrasensitive response is generated by linking low and high affinity recognition into a two-step cascade. The proposed mechanism for how our circuit is predicted to yield a cooperative T cell cytotoxic response to increasing antigen density is shown schematically in (**Fig. 2C**). Low affinity SynNotch receptors that encounter low densities of HER2 antigen would produce low amounts of CAR expressed at the T cell surface (**Fig. 2C** left). In contrast, when low affinity SynNotch receptors encounter high HER2 densities (**Fig. 2C** right) they will express higher steady-state levels of CAR. Thus, as progressively higher densities of antigen are detected, the resulting cytotoxic dose-response curve will transition as shown by the purple curves in **Fig. 2C**. Thus, increased antigen density will both increase CAR expression level, and increase the fraction of cytotoxic activation, leading to a non-linear all-or-none response (**Fig. 2C**, red line). A high killing response would only be observed when there was enough antigen to trigger sufficiently high steady state SynNotch-induced expression of the CAR.

We compared our two-step circuit strategy with the simpler approach of engineering constitutively expressed CARs with either reduced CAR expression levels or reduced CAR affinities (**Fig. S4B**). To change CAR expression level, we built a CAR tagged by a degron motif from ornithine decarboxylase (cODC) (*35, 36*). Appending the degron to our high affinity CAR construct resulted in a >10-fold reduction in CAR expression level but did not significantly change the linear killing response (**Fig. S4C**). To change CAR affinity, we replaced the high affinity scFv domain with mutant antibodies with reduced affinities for HER2 (**Fig. 1D**). Changes in CAR affinity (**Fig. 2A, Fig. S4D**) result in modest linear shifts in density dependent killing (**Fig. S4E**). Consistent with previous reports, lowering the CAR affinity can display somewhat lower toxicity against low density target cells, but also show reduced killing of high density targets (*22-24*). In summary, simply altering CAR expression levels or antigen binding affinity does not robustly increase antigen density discrimination, especially in the range required to differentiate between 0 and 3+ HER2 cells.

Given the robust behavior of the two-step HER2 antigen density sensing circuits, we tested them against well-known cancer cell lines with low and high HER2 expression levels (**Fig. 3A, Fig. S1A**). Because these circuit involves two steps, we wanted to observe the steps in T cell activation and killing using time-lapse microscopy. We collected killing time courses for the synNotch_low affinity_ →CAR_high affinty_ T cells and compared them to those of untransduced T cells or T cells constitutively expressing a high affinity CAR (**Fig. 3B and S5A**). These T cells were mixed with target cancer cells expressing either low density HER2 (PC3, 10^4.8^ Her2 molec./cell, HER2 score 1+) and high density HER2 (SKOV3, 10^7.0^ HER2 molec./cell, HER2 score 3+). For the low HER2 density PC3 cells, we observed that the constitutive high affinity CAR T cells eliminated the target cells within ∼20 hours (**Movie S1**), while neither the untransduced T cells nor the synNotch_low affinity_ → CAR_high affinity_ circuit T cells show any significant cytotoxicity over the 72 hour assay (**Fig. 3B**). Thus, the synNotch_low affinity_ → CAR_high affinity_ circuit T cells do not effectively kill the low HER2 density PC3 cells (**Movie S2)**. For the high HER2 density SKOV3 cells, we observe that constitutive high affinity CAR T cells eliminate the target cells within ∼15 hours (**Movie S1**), while the untransduced cells do not show significant cytotoxicity over the 72 hour period. In contrast, the synNotch_low affinity_ → CAR_high affinity_ circuit T cells do effectively kill the SKOV3 cells (**Movie S2**), although it exhibits a delayed activation onset and a lag in the time that it takes to achieve complete killing (72 hrs vs 24 hrs), consistent with the idea that CAR expression mediated by the synNotch pre-filter requires additional time to accumulate sufficient CAR for effective killing (**Fig. 3B**). Similar discriminatory behavior was observed against other cancer lines of varying HER2 densities (**Fig. S5A**). Direct measurement of CAR expression and T cell proliferation in co-cultures with cancer cells also showed a clear dependence on HER2 antigen density (**Fig. 3C, S5B**), as was observed against engineered K562 HER2 target cells (**Fig. 2C**). In summary, the synNotch_low affinity_ → CAR_high affinity_ T cells are able to far more cleanly discriminate between high and low density cancer cells, but the timing of circuit activation results in T cells taking a longer time to make this decision. Thus, these cells display a trade-off in which reducing the speed of the response results in a higher level of discrimination in antigen density.

**Figure 3.**
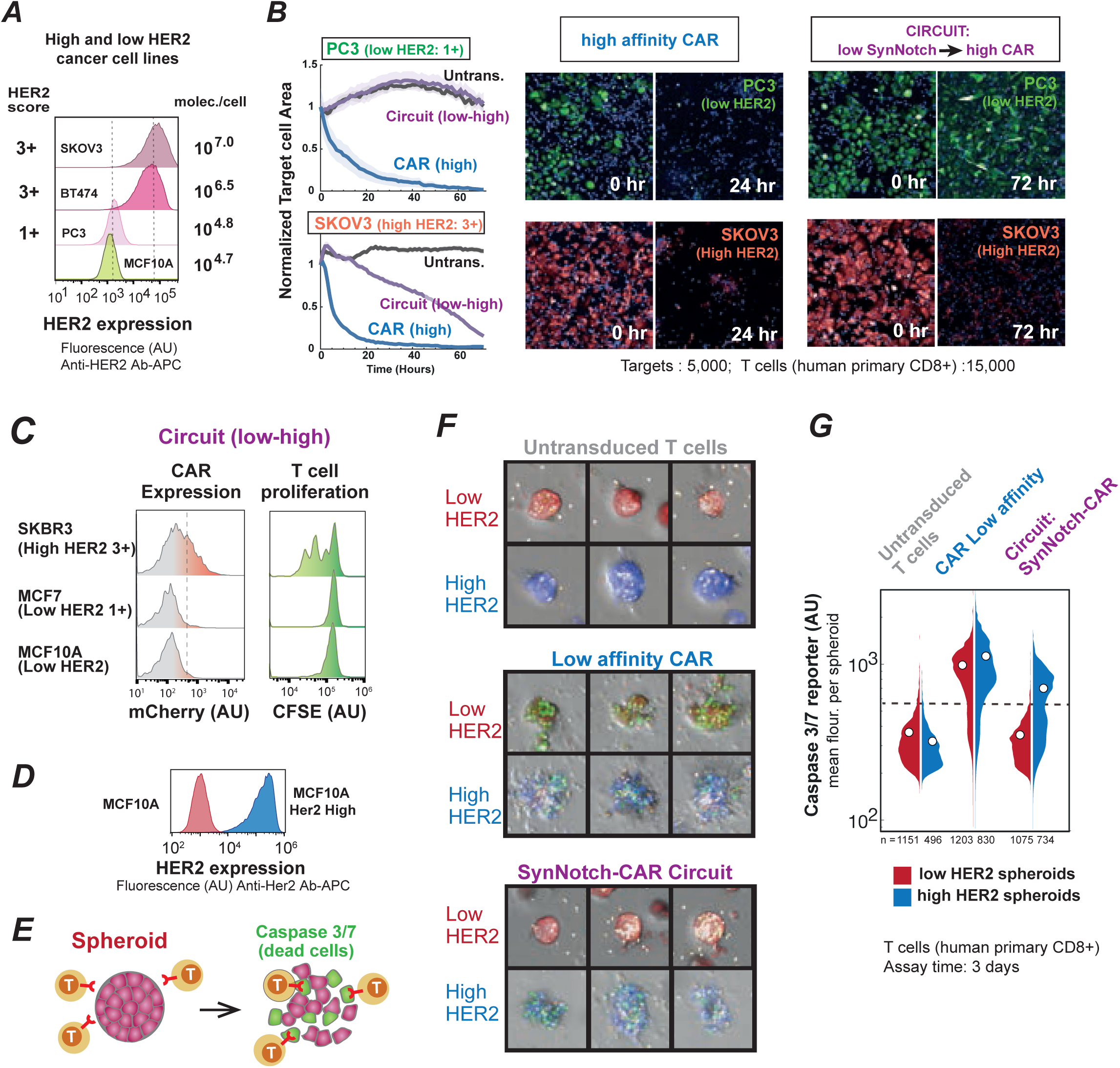
Ultrasensitive antigen density sensing T cells can efficiently discriminate between high and low density tumor cancer cell lines. **A**. FACs distributions showing the HER2 expression of low and high HER2 cancer cells. The HER2 score (as defined by ASCO-CAP scoring guidelines) is shown to the left and the average HER2 density is shown to the right. **B**. *In vitro* target cell area over time (left). Low HER2 density cancer cells (top plot), PC3 (1+ tumor line), or High HER2 density cancer cells (bottom plot), SKOV3 (3+ tumor line), were co-cultured with human primary CD8+ T cells expressing either a two-step circuit (low affinity to high affinity CAR) (purple lines) or a high affinity CAR (blue lines). Gray lines correspond to the target area in the presence of Untransduced T cells. Solid lines show the average normalized target area and the shades the standard deviation of the mean (n=3). To the right, representative images of the *in vitro* cell killing experiment. T cells are shown in blue, the low HER2 density cells in green and the high Her density cells in red (see **Fig. S5** and Supplemental **Movies S1, S2**). **C**. Representative FACS distributions for CAR expression and T cell proliferation measured for T cells expressing a low-to-high affinity recognition circuit co-cultured with normal or cancer cell lines of low and high HER2 expression. **D**. FACS distributions showing HER2 expression on MCF10A lines used to make 3D spheroids. **E**. Schematics of T cell killing assay of spheroids made of cell expressing high or low HER2. A caspase dye (shown in green) was used to track cell death. **F**. Representative images of spheroids expressing low (shown in red) or high (shown in blue) HER2 in the presence of T cell expressing either, untransduced T cells (top), a low affinity CAR (middle), or a two-step low-to-high recognition circuit (bottom). **G**. Violin plots showing the distributions of mean caspase 3/7 signal per spheroid. The distributions for the low HER2 density spheroids are shown to the left in red and the ones for the high HER2 density spheroids to the right in blue. The mean of the distribution is shown as a white circle, and the number of analyzed spheroids in each case is shown at the bottom.

In the context of immunotherapy research, we are interested on targeting solid tumors where the T cells need to interact with and infiltrate 3D multicellular aggregates. Given that it is difficult to predict the antigen density experienced by a T cell against targets organized in 3D versus 2D (*37, 38*), because in principle, a T cell that is fully surrounded by target cells in 3D, could be experiencing a higher antigen density when compared to 2D, we tested whether the antigen density sensing circuit strategy would work in a 3D culture system and in the presence of an extracellular matrix. We engineered a MCF10A line, that normally expresses low levels of HER2 (10^4.7^ Her2 molec./cell, HER2 score 0), to express high levels of HER2 (equivalent to HER2 score 3+) (**Fig. 3D**). We built 3D spheroids (*39*) using low or high HER2 MCF10A cells and embedded them in a matrix with T cells expressing either a constitutive low affinity CAR or the two-step synNotch_low affinity_ →CAR_high affinity_ circuit. Using 3D confocal microscopy and a caspase dye, we quantified the caspase fluorescence per spheroid as a proxy for cell death in each co-culture condition (**Fig. 3E, F**). Consistent to the co-cultures with cancer cells growing in two dimensions and with suspension cells (engineered K562-HER2 cells), the two-step circuit T cells discriminated high HER2 density spheroids from the low HER2 spheroids (**Fig. 3F, G**). In contrast, a low affinity anti-HER2 CAR killed low and high HER2 density spheroids indiscriminately (**Fig. 3F, G**). In all cases, the different T cells are observed to infiltrate the target cell spheroids and interact with the target cells, but they only launch a killing response (detected by caspase dye) in spheroids made of high density target cells.

To test if these density sensing T cell circuits could function *in vivo*, we evaluated their discrimination activity in dual tumor mouse models using cells with either high or low HER2 densities (below and above the desired threshold) (**Fig. 4A**). First, we implanted engineered K562 tumors on either flank of NSG mice, with a high HER2 density tumor (10^6.9^ molec./cell) on one side, and a low HER2 density tumor (10^5.2^ molec./cell) on the opposite side. After establishing the tumors, we then injected the tail vein with a mix of equal numbers of CD4+ and CD8+ primary human T cells transduced with the synNotch_low affinity_ → CAR_high affinity_ circuit (**Fig. 4B**). For comparison, we performed the same experiments with T cells constitutively expressing a low affinity anti-HER2 CAR (**Fig. S6A**). The constitutive low affinity CAR cells cleared both the low and high density tumors (no density discrimination) (**Fig. S6A**). In contrast, the synNotch_low affinity_ → CAR_high affinity_ circuit T cells show strong density discrimination (**Fig. 4B, S7A**) ╌ the high density tumors are cleared rapidly, but the low density tumors grow at similar rates as observed with untransduced T cells. Robust and consistent discrimination results were obtained in experiments with tumors grown using two other different pairs of cancer cell lines expressing high (10^7.0^ molec./cell) vs low (10^4.8^ molec./cell) levels of HER2 (**Fig. 4C**,**D, Fig. S7B, Fig. S8A**).

**Fig. 4.**
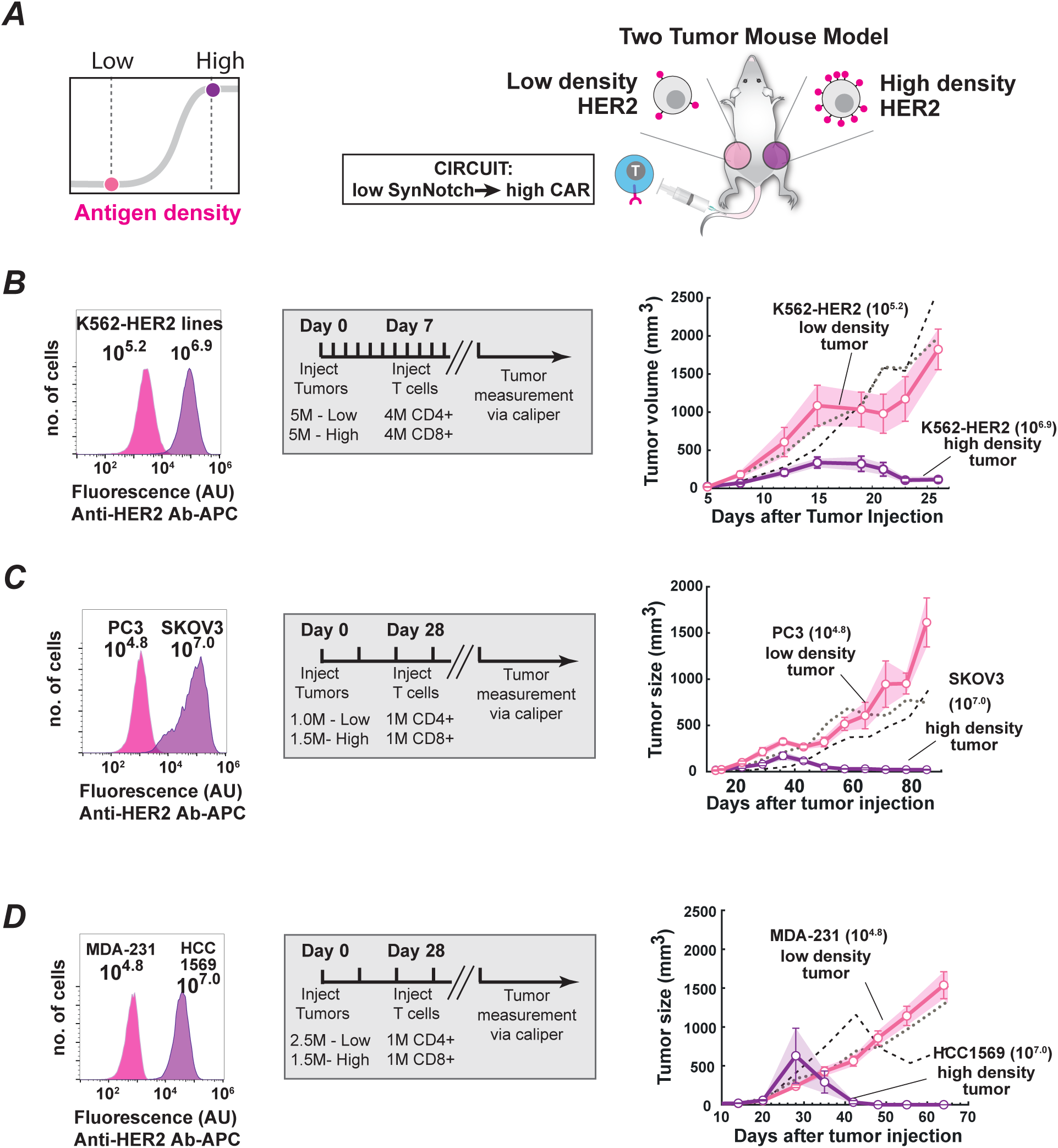
T cells expressing a low affinity SynNotch to high affinity CAR circuit show robust discrimination in several two tumor mouse models. **A**. Schematics of a two-tumor mouse model experiment to test the efficacy and safety of ultrasensitive antigen density sensing T cells. Low and high HER2 tumor cells were injected subcutaneously in the flanks of N.S.G. mice. Engineered primary human CD4+ and CD8+ T cells were injected i.v. several days after tumor injection. Tumor volume was monitored via caliper measurement over several days after tumor injection. **B-D**. FACS distributions showing the HER2 expression of cell lines utilized in the experiment. The doses and injection times for tumors and T cells are indicated in the gray box. Tumor volumes of high and low HER2 density cells after treatment with T cells expressing a two-step circuit (low affinity to high affinity CAR). The high density tumor is show in dark purple and the low density tumor in pink. The solid lines connect the means and the error bars are the standard error of the mean. **B**. Tumor volumes of engineered high and low K562 HER2 lines (n=7). The gray and black dotted lines show the low density and high density tumor volumes after treatment with untransduced T cells respectively. **C**. Tumor volumes of cancer lines, PC3 (low) and SKOV3 (high) after treatment with T cells expressing a two-step circuit (low affinity to high affinity CAR) (n=5). **D**. Tumor volumes of cancer lines, MDA-231 (low) and HCC1569 (high) after treatment with T cells expressing a two-step circuit (low affinity to high affinity CAR) (n=6). See **Figs. S6-8** for more details and individual mouse tumor volume plots.

In conclusion, our work demonstrates that a general regulatory systems design principle ╌ the use of a two-step regulatory circuit with positive feedback to generate ultrasensitive dose-response behavior ╌3 is a powerful tool to engineer T cells that can robustly differentiate between target cells with high and low antigen density expression. These two-step SynNotch to CAR circuits function well both *in vitro* and *in vivo*, and the threshold can be tuned by altering the precise affinities of the SynNotch and CAR receptors. Using this basic design, we should be able to expand the repertoire of target antigens to include other examples that are overexpressed in cancer cells compared to normal cells. Consistent with this idea, T cells engineered with a low affinity SynNotch receptor to high affinity CAR circuit, built from a toolkit of anti-EGFR binding domains (*40, 41*), efficiently discriminate engineered K562 EGFR target cells that express levels of 10^4^ from 10^6.5^ molec./cell (**Fig. S9**).

Improving our ability to effectively deploy therapeutic T cells to treat solid tumors will require several different key advances (including overcoming suppressive tumor microenvironments and improving trafficking of cells to the tumors). However, the ability to achieve ultrasensitive antigen density discrimination in engineered T cells provides a critical needed tool to treat solid cancers, in which many tumor-associated antigens are overexpressed but not absolutely unique.

## Acknowledgments

We thank Kole Roybal, Andrew Ng and Greg Allen for sharing DNA plasmids. We thank Mark Moasser for sharing tumor cell lines for *in vivo* experiments. We thank Weslley Mckeithan for assistance with microscopy data collection, Ian Eigl for assistance with biolayer interferometry data collection. We thank Aileen Li, Joseph Choe, and members of the Lim Lab for advice and helpful discussions. We thank Vu Nguyen and Greg Allen for critical reading of this manuscript.

## Funding

This work was supported by NIH grants P50GM081879 (W.A.L.), R01 CA196277 (W.A.L.), Howard Hughes Medical Institute (W.A.L.) and U54CA244438 (W.A.L.). R.H-L. is a Cancer Research Institute Irvington Fellow supported by the Cancer Research Institute. R.H-L. was a postdoctoral fellow of UC-MEXUS.

## Author contributions

R.H-L. and W.A.L. conceived the project. Experimental plan was implemented by R.H.-L., W.Y., K.C., O.C., M.L.P., Y.T. and A.G.

R.H.-L, K.C., and O.C. designed and carried out 3D culture experiments supervised by Z.J.G. and W.A.L.

R.H.-L, W.Y., Y.T., A.G. designed and carried out *in vivo* experiments supervised by W.A.L.

A.M. and K.S., designed protein expression vectors and provided scFv-GST purified protein constructs. R.H-L. analyzed all data, R.H-L., and W.A.L. prepared figures and wrote the manuscript with suggestions from all authors. W.A.L. supervised all aspects of the work.

## Competing interests

A provisional patent application has been filed by the University of California related to this work

## Data and materials availability

All data are available in the manuscript or supplementary materials. Reagents are available from the corresponding author upon reasonable request.

## Notes

### Competing Interest Statement

A provisional patent application has been filed by the University of California related to this work. Z.J.G. is an equity holder in Scribe Biosciences and Provenance Bio.

