## Supplementary Materials for "T cell circuits that sense antigen density with an ultrasensitive threshold"

**This PDF file includes:**

Materials and Methods
Figs. S1 to S9
Captions for Movies S1 and S2

**Other Supplementary Materials for this manuscript include the following:**

Movies S1 and S2

#### Materials and Methods

Gene synthesis and cloning

HER2 scFvs: Genes encoding anti-HER2 scFv variants were obtained by gene synthesis as gblocks (Integrated DNA Technologies). Gene fragments were codon optimized for expression in human cells using IDT's website tool.

EGFR scFvs and nanobodies: Similarly, genes encoding anti-EGFR scFv or nanobody variants (*1*, *2*) were obtained by gene synthesis as gblocks (Integrated DNA Technologies).
All constructs were built via in-fusion cloning (Clontech #ST0345) and sequence verified before using them.

Chimeric antigen receptor, SynNotch receptor and response element construct design

Anti-HER2 CARs were built by fusing anti-HER2 scFvs to the hinge region of the human CD8α chain and transmembrane and cytoplasmic regions of the human 4-1BB, and CD3z signaling domains. CARs were expressed under the control of a spleen focus-forming virus (SFFV) promoter. To obtain low expression levels of CARs, a degron sequence corresponding to the C-terminal region of mouse Ornithine decarboxylase (termed cODC) (EARKAIARVKRESKRIVEDLIMSCAQESAASEKISREAERLIR) was fused to the CD3z signaling domain following a $(G_4S)_3$ linker.
Anti-HER2 SynNotch receptors were built by fusing anti-HER2 ScFvs to the mouse Notch1 minimal regulatory region followed by a Gal4 DNA Binding Domain and the VP64 activation domain, as previously described (*3, 4*).
All SynNotch receptors and CAR constructs contain an N-terminal CD8a signaling peptide (MALPVTALLLPLALLLHAARP) for membrane targeting and a myc-tag (EQKLISEEDL) for determination of surface expression levels. See Morsut, Roybal et al. (*5*) for details on SynNotch receptor sequence.
The SynNotch receptors were cloned into a modified pHR'SIN:CSW vector and expressed under the control of a pGK promoter. The pHR'SIN:CSW vector was also modified to contain an inducible response element in the same plasmid. Five copies of the Gal4 DNA binding domain target sequence (GGAGCACTGTCCTCCGAACG) were cloned to a minimal CMV promoter. Inducible CAR constructs contain an N-terminal 3X-Flag Tag and C-terminal mCherry protein to determine their protein expression levels.

Engineered K562 HER2 cell lines

A series of engineered HER2 lines were derived from K562 myelogenous leukemia cells (ATCC #CCL-243). K562s were transduced with different amounts of lentivirus to stably express the extracellular and transmembrane region of human HER2 (AA 23-675) fused to a blue fluorescent protein that replaced the endogenous intracellular region. The HER2-BFP construct was expressed under the control of the SFFV promoter. Cells were sorted on an Aria Fusion cell sorter (BD Biosciences) on the basis of HER2 expression and subsequently expanded and frozen at low passage. HER2 levels in each target cell population were determined by staining the cells with anti-HER2 APC (Biolegend 324407); see details below. Overexpression of HER2 is consistent with the amplified levels found in +3, +2 and +1 tumors as scored by ASCO-CAP scoring guidelines (Fig. 1B, Fig. S1A). All engineered K562 cell lines were subcultured in IMDM media supplemented with 10% FBS and gentamicin.

Engineered MCF10A HER2 cell lines

A set of HER2 overexpression lines were derived from MCF10As. MCF10A is a non-tumorigenic epithelial cell line capable of three-dimensional growth in matrigel or collagen. MCF10As were transduced with different amounts of lentivirus to stably express the extracellular and transmembrane region of human HER2 (AA 23-675) fused to a blue fluorescent protein that replaced the endogenous intracellular region. The HER2-BFP construct was expressed under the control of the SFFV promoter. Cells were sorted on an Aria Fusion cell sorter (BD Biosciences) on the basis of HER2 expression and subsequently expanded and frozen at low passage.

Engineered EGFR cell lines

Similarly, a series of EGFR amplified lines was obtained using a construct that expresses the extracellular region (AA 1-645) of EGFR and transmembrane region of PDGFR (AA 512-561). All cell lines were stained and sorted for expression of transgenes using an anti-EGFR BV786 (BD Bioscience 742606) antibody. Engineered K562-EGFR cell lines were subcultured in IMDM media supplemented with 10% FBS and gentamicin.

Cancer cell lines culturing protocols

All cancer cell lines used in this study were purchased from the indicated vendors. Cells were cultured to confluence in the indicated media supplemented with 10% FBS. At each passage, cells were washed with PBS (at 37 °C) and TrypLE (ThermoFisher Scientific 12604021) was added to detach the cells from the flask surface. Flasks were incubated at 37 °C until the cells detached, typically 5 to 10 min. Fresh culture medium was added to quench the TrypLE and cells were resuspended and plated in new flasks and in fresh culture medium. PC3 cells (ATCC CRL-1435) were cultured in F-12K medium, SKOV3 cells (ATCC CRL-HTB77) in McCoy's 5a medium, MCF7 cells (ATCC CRL-HTB22) in DMEM medium, BT474 cells (ATCC CRL-HTB20) in RPMI medium and MCF10-A (ATCC CRL-10317) in DMEM/F-12 medium supplemented with 5% horse serum, cholera toxin to a final concentration of 1 ng/mL; human insulin to a final concentration of 10 ug/mL; epidermal growth factor to a final concentration of 10 ng/mL; and hydrocortisone to a final concentration of 0.5 ug/mL. MDA-231-M1 and HCC1569-M1 cell lines were kind gifts of the Moasser Lab (UCSF).

Determination of protein copy number per cell

Average Antigen density per target cell was determined by quantitative flow cytometry. For each of the target antigens or receptors, $1x10^5$ cells of each population were stained with the desired antibody: Anti-HER2 APC (Biolegend 324407), Anti-EGFR BV786 (BD Biosciences 742606) or Anti-myc Alexa 647 (CellSignaling 2233S) antibody for 30 min on ice (n=3). Cells were washed twice with PBS and resuspended in PBS for analysis in an Attune NxT Flow Cytometer. The geometric mean of each target population was determined after gating the cells by their size (side scatter and forward scatter region) and selecting the full width at half maximum (FWHM) of the population in the corresponding fluorescent channel. A standard curve was built using Quantum Simply Cellular anti-Mouse IgG beads (Bang Laboratories 815) stained with the same antibody than the target cells. For each cell line, the number of molecules per cell was determined using the standard curve and the geometric mean of each target population. Similarly, to determine the expression amounts of inducible CAR expression, mCherry flow cytometer calibration beads (Takara Bio 632595) were used.

Primary human T Cell isolation and culture

Primary CD4+ and CD8+ T cells were isolated from blood of anonymous donors by negative selection (STEMCELL Technologies #15062 and #15063). T cells were cryopreserved in RPMI-1640 (UCSF cell culture core) with 20% human AB serum (Valley Biomedical, #HP1022) and 10% DMSO. T cells were cultured in human T cell medium consisting of X-VIVO 15 (Lonza #04-418Q), 5% Human AB serum, and 10 mM neutralized N-acetyl L-Cysteine (Sigma-Aldrich #A9165) supplemented with 30 units/mL IL-2 (NCI BRB Preclinical Repository) for all experiments.

Lentiviral transduction of human T cells

Pantropic VSV-G pseudotyped lentivirus was produced by transfecting Lenti-X 293T cells (Clontech #11131D) with a pHR'SIN:CSW transgene expression vector and the viral packaging plasmids pCMVdR8.91 and pMD2.G using Fugene HD (Promega #E2312). Primary T cells were thawed and after 24 hr in culture, were stimulated with Human T-Activator CD3/CD28 Dynabeads (Life Technologies #11131D) at a 1:1 cell:bead ratio. After 48 hr, viral supernatant was harvested and added to primary T cells. T cells were exposed to the virus for 24 hr. At day 5 after T cell stimulation, the Dynabeads were removed. T cells were stained and sorted on an Aria Fusion cell sorter (BD Biosciences) on the basis of myc stain to obtain homogenous receptor expression levels. For the SynNotch to CAR circuits, T cells expressing CAR were removed by cell sorting. T cells were expanded for at least 9 days when they were rested and could be used for killing assays.

ScFv protein expression and purification

For binding affinity measurements, genes encoding the Anti-HER2 scFv fragments used to build the SynNotch and CAR constructs (including the N-terminal CD8a signaling peptide and the myc-tag) were cloned into a pEBB-GST vector. The pEBB-GST vector was constructed by inserting a GST from *Schistosoma japonicum* transcarboxylase into the polylinker site of the expression vector pEBB. These fusion constructs are under the control of an EF1α promoter. Expression vectors were used to transiently transfect HEK 293 cells. All proteins were purified from filtered cell supernatants according to the instructions provided with the glutathione-sepharose 4B resin (GE Healthcare). Proteins were dialyzed against PBS after the glutathione elution and flash frozen in small aliquots.

Biolayer interferometry

GST-tagged scFvs were immobilized to an anti-GST capture sensor tip (FortéBio) using an Octet RED384 (FortéBio). The sensors were preconditioned according to the FortéBio protocol and then dipped into a series of concentrations of label free HER2 (Acro byosystems; HE2-H5212) to measure association before being dipped into a well containing only running buffer composed of 1X PBS, 0.05% Tween 20, 0.3 % bovine serum albumin to measure dissociation.
Data were reference subtracted and fit to a 1:1 binding model using Octet Data Analysis Software v11.1 (FortéBio).

*In vitro* T cell cytotoxicity assessment

CD8+ Primary human T cells expressing either constitutive anti-HER2 CAR or two-step SynNotch-CAR circuits were co-cultured for 72 hr with targets expressing different HER2 densities in complete human T cell medium and placed at 37 °C, 5% $CO_2$ incubator. For all *in vitro* T cell killing assays against engineered K562-HER2 cells T cells were stained with celltrace CFSE dye (Thermo Fisher Scientific C34554) and co-cultured in round bottom 96-well tissue culture plates at the indicated effector to target ratios. The cells were centrifuged for 1 min at 400 x g to favor effector to target interactions, and the cultures were analyzed at 72 Hrs. for specific lysis of target tumor cells, T cell proliferation and CAR expression by flow cytometry. T cells were identified by the celltrace dye and target cells were identified by size, see Fig. S4A. The level of specific lysis of target cancer cells was determined by comparing the number of target cells alive in the culture compared to treatment with untransduced T cell controls. Cell death was monitored by a live-dead cell stain and by shifting of the target cells out of the side scatter and forward scatter region normally populated by the target cells. All flow cytometry was performed using BD LSR II or Attune NxT Flow Cytometers and the analysis was performed in FlowJo software (TreeStar) and Matlab.

Imaging of T cell cytotoxicity- 2D cultures

For all *in vitro* T cell killing of HER2 expressing cancer lines, $5x10^3$ target cells were stained with a celltrace dye, cultured overnight in a flat bottom 384-well tissue culture plate in their indicated medium and placed at 37 °C, 5% $CO_2$ incubator. After 1 day, $1.5 \times 10^4$ T cells were stained and added to the flat bottom 384-well tissue culture plate and the co-cultures were imaged every hour for 3 days. Two fields per well were imaged using the 20x objective on a PerkinElmer Opera Phenix High Content Screening System and the images were analyzed using the associated Harmony Office Software. Data was summarized as the sum of the normalized area occupied by target cells and presented as mean ± SEM.

*In vitro* T cell proliferation and CAR expression with cancer cell lines

For *in vitro* assessment of T cell proliferation and CAR expression with cancer lines, $10x10^3$ target cells were cultured overnight in a flat bottom 96-well tissue culture plate in their indicated medium and placed at 37 °C, 5% $CO_2$ incubator. After 1 day, $2.5 \times 10^4$ T cells were stained, following the manufacturer protocol, and added to the flat bottom 96-well tissue culture plate. The plates were centrifuged for 1 min at 400 x g to favor effector to target interactions, and cultures were analyzed at 72 hr. T cell proliferation and CAR expression were determined by flow cytometry.

*In vitro* assessment of T Cell Cytotoxicity – 3D cultures

To assess the killing activity of constitutive anti-HER2 CAR T cells or two-step $\text{SynNotch}_{\text{low affinity}}$ → $\text{CAR}_{\text{high affinity}}$ circuit T cells in the context of a three-dimensional tissue, we created aggregates of MCF10A cells using agarose microwells, as previously described (*6*). Briefly, photolithography was used to create a master wafer with SU-8 microwells of 120 µm diameter, 100 µm height. PDMS micropillar stamps were created from this master by pouring Sylgard 184 (Dow Corning, 10:1 base:crosslinker ratio) over the wafer and baking at 60˚C for >3 hours. The PDMS stamps were placed on top of molten agarose (3% weight/volume in PBS (calcium and magnesium free), AllStar Scientific) in 6-well polystyrene plates (Thermo Fisher). After the agarose cooled, the PDMS stamps were peeled off, leaving behind 120 µm agarose microwells. MCF10A cells were re-suspended in PBS + 0.4% EDTA at a density of $1 \times 10^6$ cells/mL and added to the well plate containing the agarose microwells. The MCF10As were centrifuged into the agarose microwells at 160 x g for four minutes, the plate was rotated 180 degrees, and then centrifuged a second time. Excess cells that did not occupy the microwells were removed by washing with PBS + EDTA followed by washing with MCF10A media. After overnight culture, the MCF10As in each microwell had coalesced to form spheroidal cell aggregates.

MCF10A aggregates were removed from the agarose microwells, sedimented with pulsed centrifugation to 20G, and then resuspended in ice-cold Matrigel containing 1 x $10^6$ T cells/mL. The Matrigel + T cell + MCF10A aggregate mixture was plated into Matrigel-coated wells of a glass bottomed 24-well plate (Cellvis). Each well was given Human T cell media containing 30 units/mL IL-2, 20 ng/mL EGF and 2 drops/mL of CellEvent Caspase 3/7 Green Dye (Invitrogen). The Caspase 3/7 dye served as a readout of cell death.
Co-cultures were imaged after 3 days. Several fields and 15 planes, 50 µm apart, per well were imaged using the 5x objective on a PerkinElmer Opera Phenix High Content Screening System and the images were analyzed using the associated Harmony Office Software. Data was summarized as the mean caspase fluorescence intensity per spheroid and presented as violin plots.

Statistical analysis and curve fitting

Data is presented as means ± standard error of the mean (SEM) or means ± standard deviations (SD) as indicated in the figure legends. The target cell killing data for the SynNotch to CAR circuits was fitted to a four parameter Hill equation using the curve fitting toolbox in Matlab (see **Fig. S3B** for details of the equation an parameters).

*In vivo* mouse models

All mouse experimental procedures were conducted according to Institutional Animal Care and Use Committee (IACUC)–approved protocols. Female NSG mice were obtained from UCSF breeding core. To evaluate the safety and efficacy of the $\text{SynNotch}_{\text{low affinity}} \rightarrow \text{CAR}_{\text{high affinity}}$ circuit, 6- to 12- week-old animals were inoculated with tumor cells in PBS solution, subcutaneously in the right (high density) and left (low density) flanks, respectively. Single dose treatments consisting of sorted and rested CD4+ and CD8+ engineered or the matched number of untransduced T cells were administered intravenously via tail vein in 100 µL of PBS at day 7 after tumor injection. Tumor volumes were monitored two times a week via caliper measurements until predetermined IACUC-approved endpoint (hunching, neurological impairments such as circling, ataxia, paralysis, limping, head tilt, balance problems, seizures, tumor volume burden) was reached (n = 5 to 7 mice per group). For experiments with cancer cell lines, cell suspensions in PBS were mixed 1:1 with Matrigel (Corning) and 50 µL were inoculated subcutaneously in the right (high density) and left (low density) flanks, respectively. T cells were administered intravenously via tail vein in 100 µL of PBS at day 28 after tumor injection. Tumor volumes were monitored once a week via caliper measurements until predetermined IACUC-approved endpoint (hunching, neurological impairments such as circling, ataxia, paralysis, limping, head tilt, balance problems, seizures, tumor volume burden) was reached (n = 5 to 7 mice per group).

#### Fig. S1. Determination of antigen density and receptor expression from fluorescence intensity

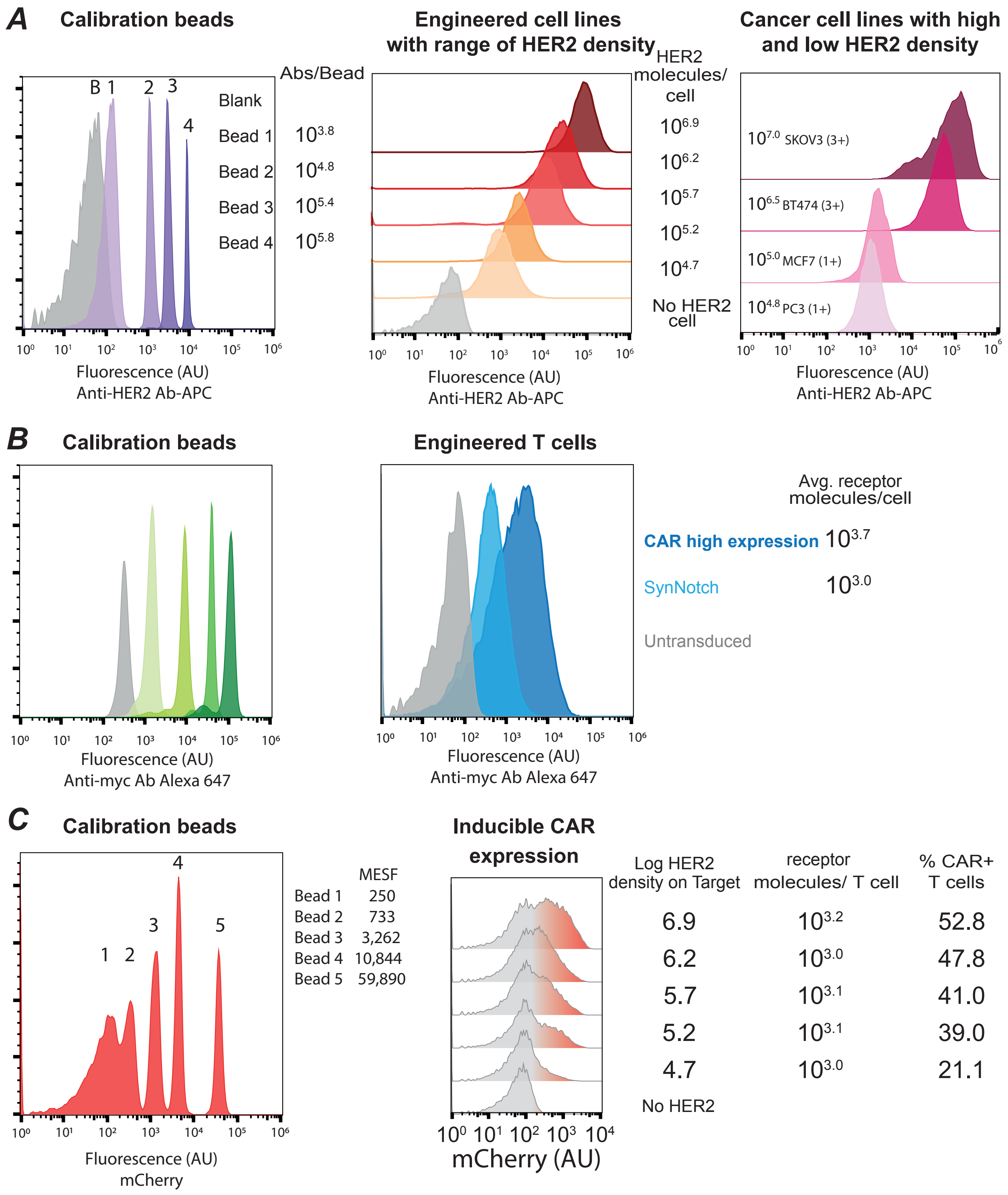

**Figure S1. Determination of antigen density and receptor expression from fluorescence intensity.**

Antigen density and receptor expression were determined by quantitative flow cytometry. **A.** left: Representative flow cytometry histograms showing the fluorescence intensity of Quantum Simply Cellular anti-Mouse IgG beads (Bang Laboratories 815) stained with anti-HER2 APC antibody. The manufactured antibody binding capacity of each bead population is indicated to the right in top panel. Center: Representative flow cytometry histograms of engineered K562 HER2-BFP cell lines stained with anti-HER2 APC antibody. The geometric mean of each population and a calibration curve built from data shown in the left panel was used to determine the number of HER2 molecules per cell in each population. Right: Representative flow cytometry histograms showing the fluorescence intensity of cancer cell lines expressing a range of HER2 densities. The density of HER2 molecules/cell and their classification as scored by ASCO-CAP scoring guidelines is shown. **B.** Left: Similar to A for beads stained with anti-myc Alexa 647. Center: Engineered CD8+ human T cells expressing either a constitutive CAR or a SynNotch receptor were stained with anti-myc Alexa 647. The number of receptors per T cell populations was determined as described above. **C.** Left: Representative flow cytometry histograms of beads showing fluorescence intensity equivalent to the indicated number of soluble mCherry molecules (MESF). Center: Representative histograms showing the fluorescence intensity of CD8+ human T cells co-cultured with K562 HER2-BFP target cells for 3 days. The geometric mean of the positive population and the corresponding calibration curve was used to determine the number of induced CAR molecules per cell in each population. The percentage of CAR positive cells was determined using the population comparison platform in FlowJo V10 and it is reported as % SE Dymax. Briefly, it normalizes the data to a unit scale to protect against outliers, and factors in the distribution of the data.

### Fig. S2. Protein sequences for anti-HER2 scFvs and their binding affinities

## *A*

| | 1 | 11 | 21 | 31 | 41 |
|---|---|---|---|---|---|
| 4D5_WT_Highest | D I Q M T Q S P S S | L S A S V G D R V T | I T C R A S Q D V N | T A V A W Y Q Q K P | G K A P K L L I Y S |
| 4D5_Medium | D I Q M T Q S P S S | L S A S V G D R V T | I T C R A S Q D V N | T A V A W Y Q Q K P | G K A P K L L I Y S |
| 4D5_High | D I Q M T Q S P S S | L S A S V G D R V T | I T C R A S Q D V N | T A V A W Y Q Q K P | G K A P K L L I Y S |
| 4D5_Low | D I Q M T Q S P S S | L S A S V G D R V T | I T C R A S Q D V N | T A V A W Y Q Q K P | G K A P K L L I Y S |

| | 51 | 61 | 71 | 81 | 91 |
|---|---|---|---|---|---|
| 4D5_WT_Highest | A S F L Y S G V P S | R F S G S R S G T D | F T L T I S S L Q P | E D F A T Y Y C Q Q | H Y T T P P T F G Q |
| 4D5_Medium | A S F L E S G V P S | R F S G S R S G T D | F T L T I S S L Q P | E D F A T Y Y C Q Q | H Y T T P P T F G Q |
| 4D5_High | A S F L E S G V P S | R F S G S R S G T D | F T L T I S S L Q P | E D F A T Y Y C Q Q | H Y T T P P T F G Q |
| 4D5_Low | A S F L E S G V P S | R F S G S G S G T D | F T L T I S S L Q P | E D F A T Y Y C Q Q | H Y T T P P T F G Q |

| | 101 | 111 | 121 | 131 | 141 |
|---|---|---|---|---|---|
| 4D5_WT_Highest | G T K V E I K R T G | S T S G S G K P G S | G E G S E V Q L V E | S G G G L V Q P G G | S L R L S C A A S G |
| 4D5_Medium | G T K V E I K R T G | S T S G S G K P G S | G E G S E V Q L V E | S G G G L V Q P G G | S L R L S C A A S G |
| 4D5_High | G V K V E I K R T G | S T S G S G K P G S | G E G S E V Q L V E | S G G G L V Q P G G | S L R L S C A A S G |
| 4D5_Low | G V K V E I K R T G | S T S G S G K P G S | G E G S E V Q L V E | S G G G L V Q P G G | S L R L S C A A S G |

| | 151 | 161 | 171 | 181 | 191 |
|---|---|---|---|---|---|
| 4D5_WT_Highest | F N I K D T Y I H W | V R Q A P G K G L E | W V A R I Y P T N G | Y T R Y A D S V K G | R F T I S A D T S K |
| 4D5_Medium | F N I K D T Y I H W | V R Q A P G K G L E | W V A R I Y P T N G | Y T R Y A D S V K G | R F T I S A D T S K |
| 4D5_High | F N I K D T Y I H W | V R Q A P G K G L E | W V A R I Y P T N G | Y T R Y A D S V K G | R F T I S A D T S K |
| 4D5_Low | F N I K D T Y I H W | V R Q A P G K G L E | W V A R I Y P T N G | Y T R Y A D S V K G | R F T I S A D T S K |

| | 201 | 211 | 221 | 231 | 241 |
|---|---|---|---|---|---|
| 4D5_WT_Highest | N T A Y L Q M N S L | R A E D T A V Y Y C | S R W G G D G F Y A | M D V W G Q G T L V | T V S S G S |
| 4D5_Medium | N T A Y L Q M N S L | R A E D T A V Y Y C | S R W G G D G F Y A | M D V W G Q G T L V | T V S S G S |
| 4D5_High | N T A Y L Q M N S L | R A E D T A V Y Y C | S R W G G D G F Y A | M D V W G Q G T L V | T V S S G S |
| 4D5_Low | N T A Y L Q M N S L | R A E D T A V Y Y C | S R W G G D G F Y A | M D V W G Q G T L V | T V S S G S |

## *B*

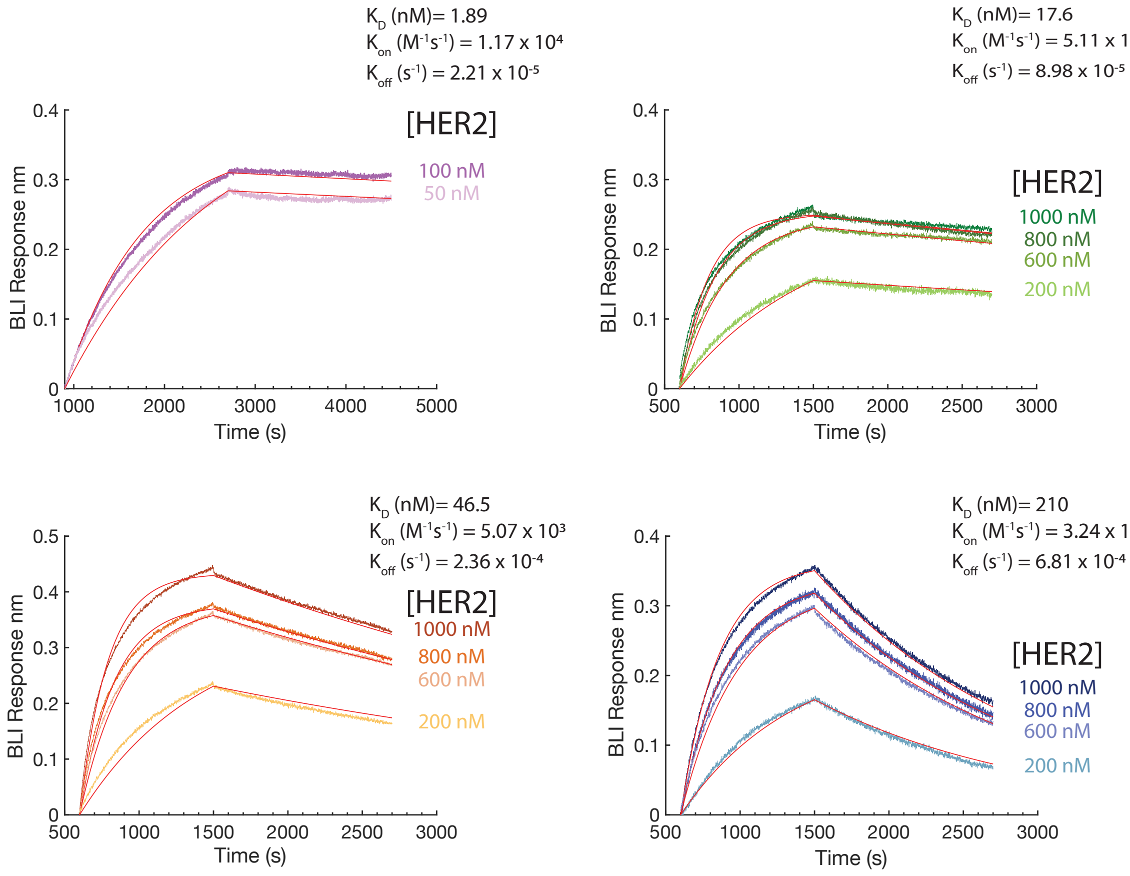

**Figure S2. Protein sequences for anti-HER2 scFvs and their binding affinities**

**A.** Sequence alignment of scFvs derived from Anti-HER2 4D5 antibody. The WT sequence is labeled as 4D5-WT-Highest and the residues that were mutated to create each variant are indicated in red. **B.** Biolayer interferometry sensograms showing the binding kinetics for human HER2 and immobilized Anti-HER2 scFv variants. Data are shown as colored lines and the best fit for data to a 1:1 binding model is shown in red. The binding affinities, association and dissociation constants that resulted from the fit are show. HER2 concentrations utilized for binding affinity measurements are indicated**.**

#### Fig. S3 Effects of receptor affinity and T cell dosage on two-step circuit function

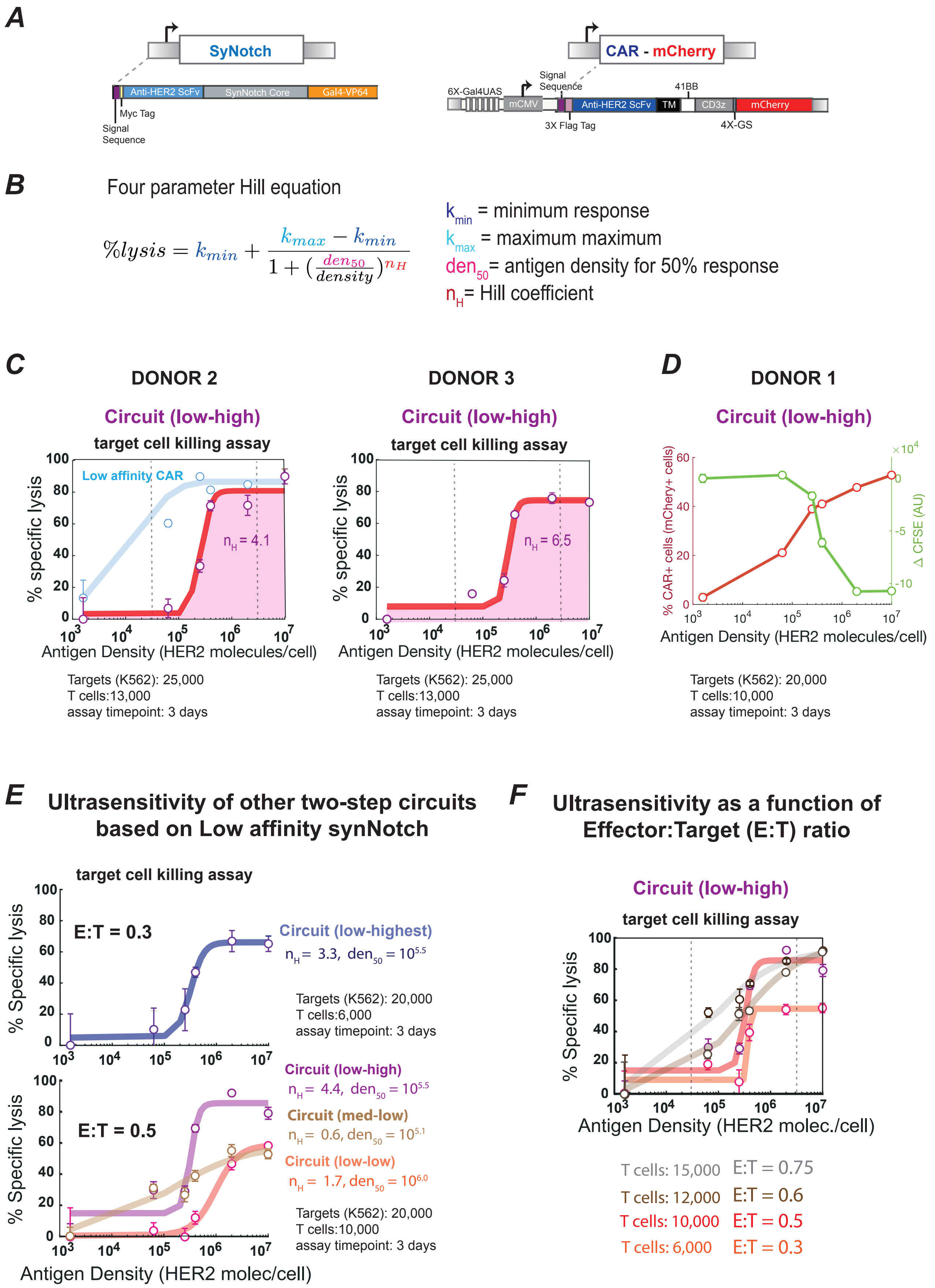

**Figure S3. Effects of receptor affinity and T cell dosage on two-step circuit function.**

**A.** Construct design for a two-step SynNotch to CAR circuit. The main domains are indicated and color coded (see methods for details). **B.** Four parameter Hill equation utilized to fit the killing response curves as a function of antigen density of two-step circuits tested in this study. The four parameters are color coded and indicated to the right of the equation. **C.** Target cell killing response curve for T cells expressing low affinity SynNotch to high affinity CAR circuit from other two different donors. For donor 2 as a comparison the curve for the constitutive expression of a low affinity CAR is shown. For the circuits transparent lines are fits to a hill equation, the hill coefficient for each curve is indicated. For constitutive CARs, transparent lines are drawn based on inspection. **D.** %CAR+ cells and dilution of cell trace marker CFSE to assess the CAR expression and T cell proliferation as a function of target cell HER2 density (at 3 days) for T cells expressing a low-to-high affinity recognition circuit (See **Fig. 2B**). **E.** Target cell killing response curves for T cells expressing different two-step circuits. A low affinity SynNotch receptor was used in these circuits. Different affinity CAR receptors were tested. The low affinity SynNotch to low affinity CAR circuit showed a higher antigen threshold than other designs. The hill coefficient ($n_H$) and antigen density for the half maximal activity ($den_{50}$) values are indicated for each killing curve. **F.** Target cell killing response curves for T cells expressing low affinity SynNotch to high affinity CAR circuit at different effector to target (E:T) ratios. Ultrasensitivity is best observed at low E:T ratios.

### Figure S4. Altering CAR expression level or affinity yields modest linear changes in antigen density sensitivity.

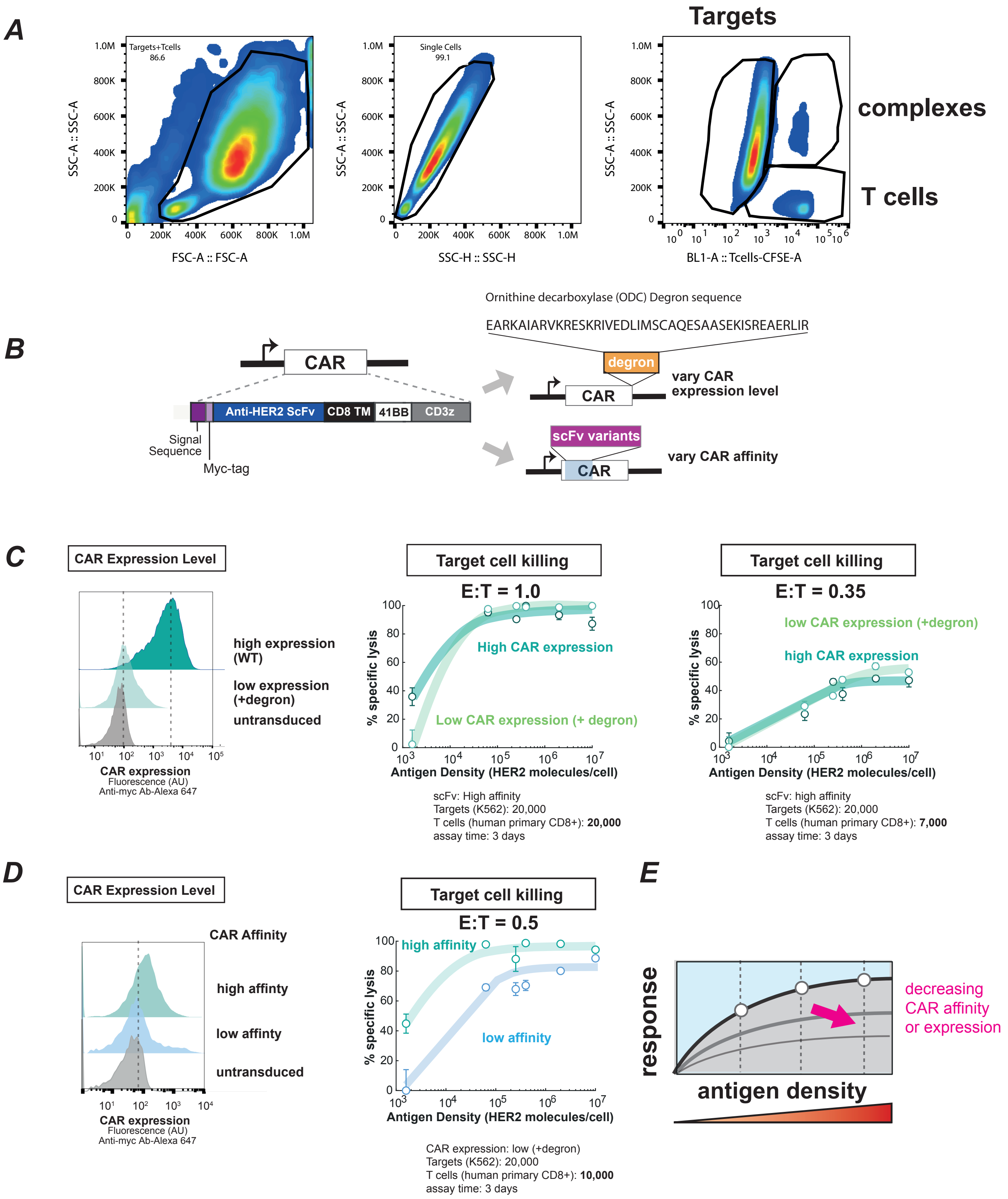

**Figure S4. Altering CAR expression level or affinity yields modest linear changes in antigen density sensitivity.**

**A.** Details on gating scheme utilized to analyze killing assays by flow cytometry. Samples were first gated using a live-dead cell stain dye (not shown), then using forward and side scattering profiles to select single cells and finally using the CFSE celltrace fluorescence to separate T cells from K562-HER2 targets. T cell-Target complexes were excluded from the analysis. **B**. Construct Design of anti-HER2 CARs used in this study. We obtained T cells expressing varying levels of receptor by fusing a degron tag to the CAR. We altered CAR affinity by changing the scFv domain. The degron sequence corresponds to the C-terminal region of mouse Ornithine decarboxylase (termed cODC) (EARKAIARVKRESKRIVEDLIMSCAQESAASEKISREAERLIR) which was fused to the original CAR constructs to reduce CAR expression. **C.** Effect of changing CAR expression levels on antigen density dependent cell killing (here we use the high affinity scFv). Representative FACs histograms for CAR expression distribution in human primary CD8+ T cells, with and without degron tag. Plots on right shows antigen density dependence of target cell killing at two different T cell doses. **D.** Effect of changing CAR affinity on antigen density dependent cell killing (here we fix low CAR expression - with a degron tag). FACS plots on left show CAR expression distribution of each affinity CAR in human primary CD8+ T cells. Plot on right shows antigen density dependence of target cell killing. Transparent lines are drawn based on inspection. The percentage of specific lysis was determined using quantitative flow cytometry by counting the number of target cells after 3 days relative to a co-culture in the presence of untransduced T cells. **E.** Schematics of effects on changing CAR affinity or CAR expression on antigen density dependent T cell killing response. Decreasing CAR affinity or expression levels leads to linear changes in antigen density response curves.

**Fig. S5. Low affinity SynNotch to high affinity CAR T cells show antigen density activity against several HER2 positive cancer lines in vitro.**

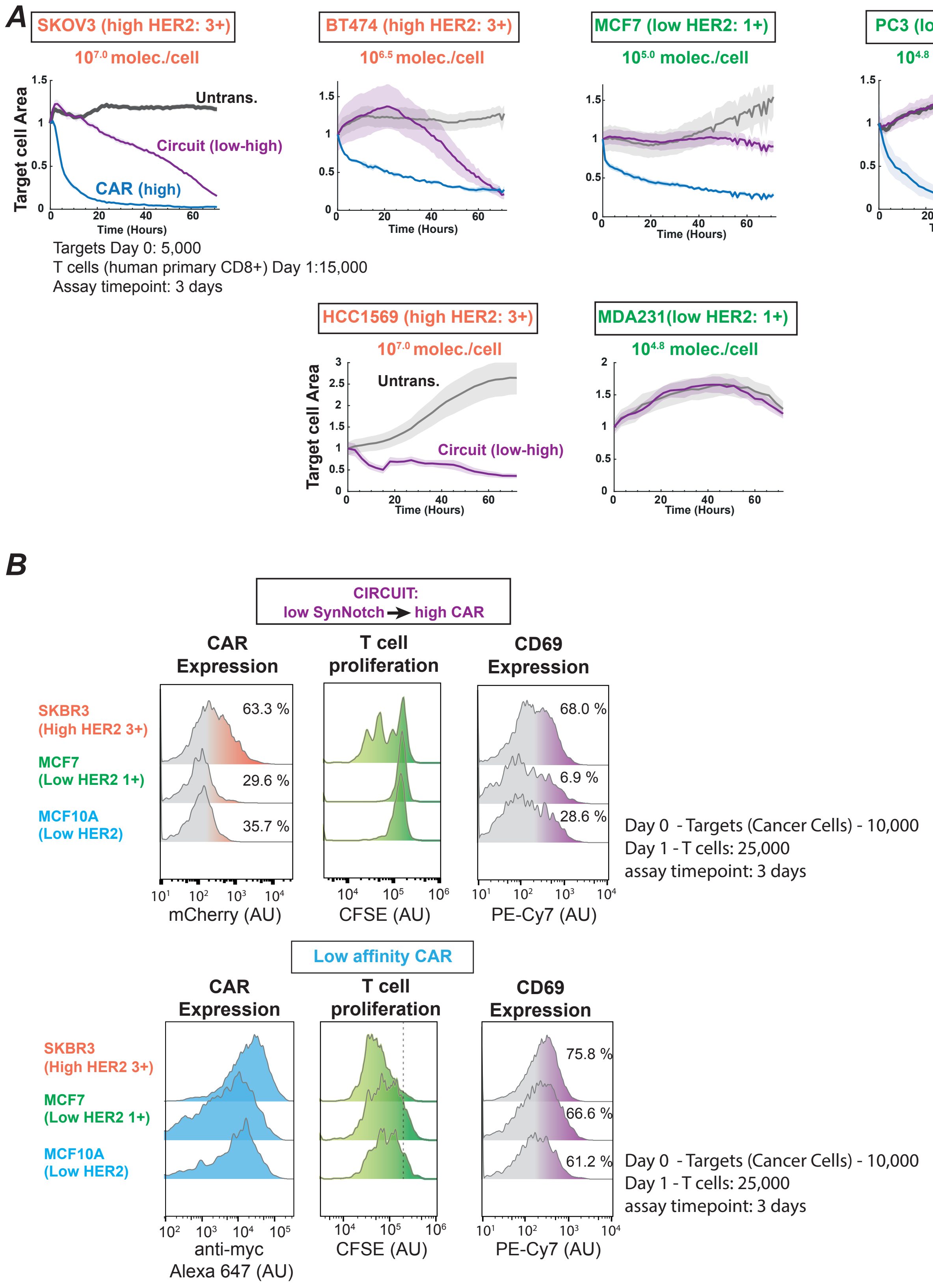

**Figure S5. Low affinity SynNotch to high affinity CAR T cells show antigen density activity against several HER2 positive cancer cell lines *in vitro*.**

**A.** *In vitro* target cell area over time. Cancer lines with several HER2 densities were co-cultured with human primary CD8+ T cells expressing either a two-step circuit (low affinity to high affinity CAR) (purple lines) or a high affinity CAR (blue lines). Gray lines correspond to the target cell area in the presence of Untransduced T cells. Solid lines show the average target area and shaded areas show the standard deviation (n=3). The mean HER2 density and classification is indicated for each cancer line. **B.** Representative FACS plots of CAR expression, T cell proliferation and CD69 expression for T cells co-cultured with cancer cell lines expressing high and low HER2 densities. The histograms in the top row correspond to the two-step SynNotch to CAR circuit T cells. The histograms in the bottom row correspond to a constitutive low affinity CAR. The E:T and assay time point are indicated.

**Fig. S6. T cells expressing a constitutive low affinity CAR do not discriminate tumors expressing high and low HER2 levels**

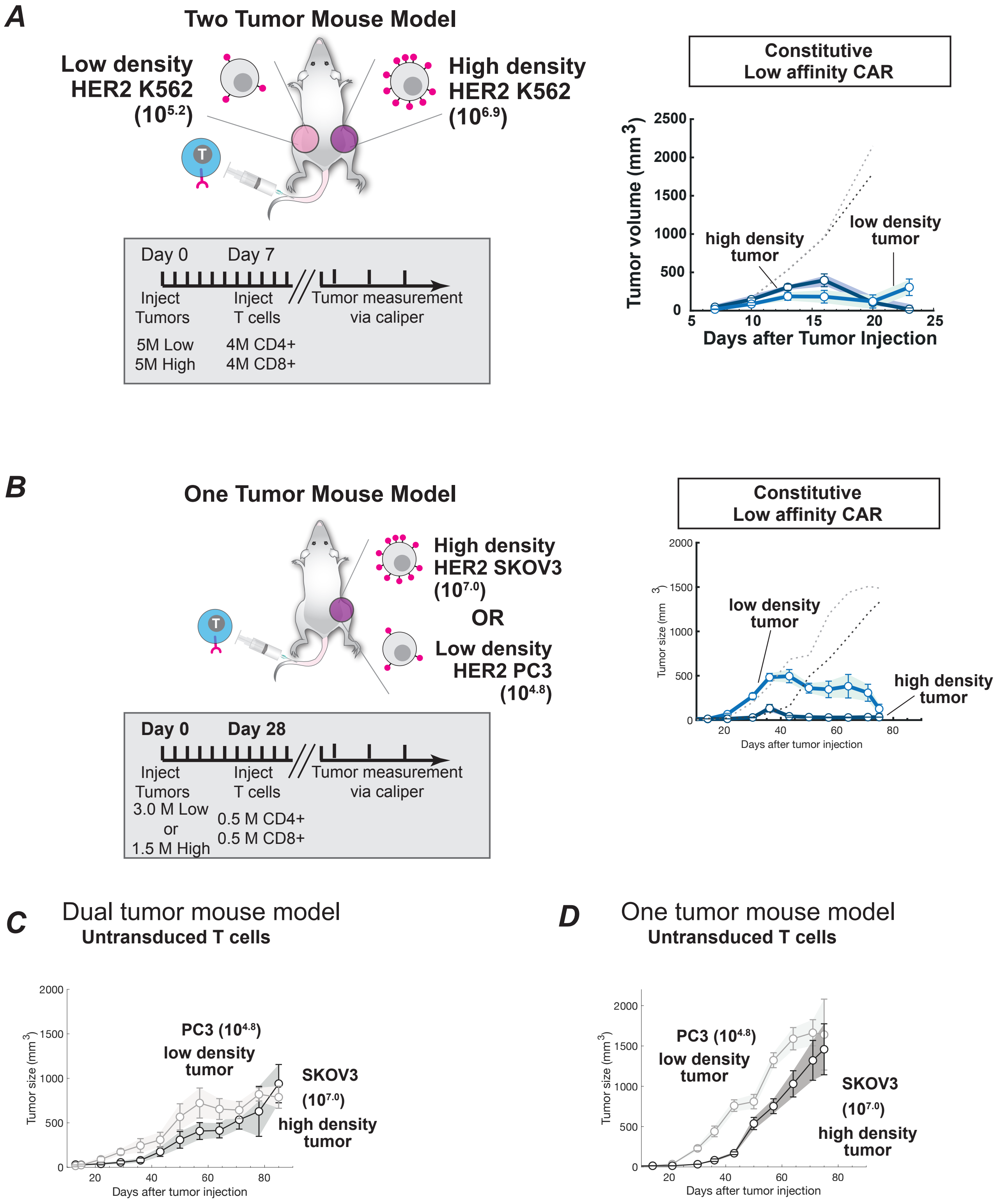

**Figure S6. T cells expressing a constitutive low affinity CAR do not discriminate tumor expressing high and low HER2 levels.**

**A.** Schematics of a two-tumor mouse model experiment. Low and high engineered K562-HER2 tumor cells were injected subcutaneously in the flanks of N.S.G. mice. Engineered primary human CD4+ and CD8+ T cells were injected i.v. 7 days after tumor injection. Tumor volume was monitored via caliper measurement over several days after tumor injection. The dark blue line corresponds to the high HER2 K562 tumor whereas the light blue lines corresponds to the low HER2 K562 tumor after treatment with constitutive low affinity CAR (n=8). The gray and black dotted lines show the low density and high density mean tumor volumes after treatment with untransduced T cells (n=8). **B.** Schematics of a one-tumor mouse model experiment. Low or high HER2 tumor cells were injected subcutaneously in the flanks of N.S.G. mice. Engineered primary human CD4+ and CD8+ T cells were injected i.v. 28 days after tumor injection. Tumor volume was monitored via caliper measurement over several days after tumor injection. The dark blue line corresponds to the high (SKOV3, n=5) tumor whereas the light blue line corresponds to the low (PC3, n=6) tumor after treatment with constitutive low affinity CAR. The gray and black dotted lines show the low density (n=6) and high density (n=6) mean tumor volumes after treatment with untransduced T cells respectively. **C.** The gray and black lines show the low density and high density mean tumor volumes (n=5), respectively, after treatment with untransduced T cells for the experiment shown in **Fig. 4C**,**D.** The gray and black lines show the low (n=6) density and high (n=6) density mean tumor volumes, respectively, after treatment with untransduced T cells for the experiment shown in **Fig. S6B**.

**Fig. S7. Tumor volume measurements for individual mice treated with T cells expressing low affinity SynNotch to high affinity CAR circuit.**

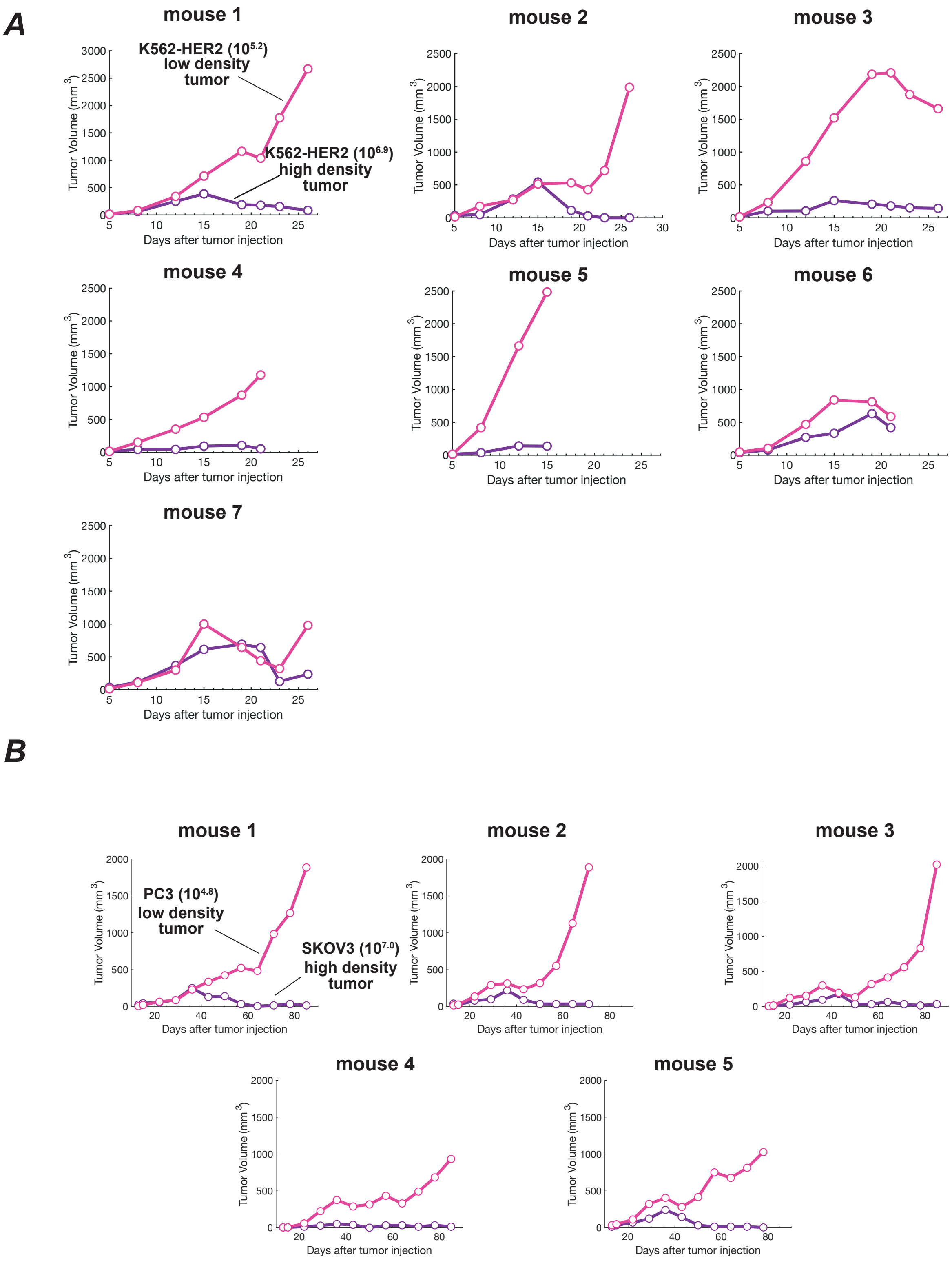

**Figure S7. Tumor volume measurements for individual mice treated with T cells expressing low affinity SynNotch to high affinity CAR circuit.**

**A.** Tumor volume data for individual mice treated with T cells expressing a low affinity SynNotch to high affinity CAR circuit. The dark purple lines correspond to the high HER2 K562 tumor whereas the light pink lines correspond to the low HER2 K562 tumor (See **Fig. 4B**). **B.** Tumor volume data for individual mice treated with T cells expressing a low affinity SynNotch to high affinity CAR circuit. The dark purple lines correspond to the high (SKOV3) tumor whereas the light pink lines correspond to the low (PC3) tumor (See **Fig. 4C**).

**Fig. S8. Tumor volume measurements for individual mice treated with T cells expressing low affinity SynNotch to high affinity CAR circuit, related to Fig. 4D.**

***A***

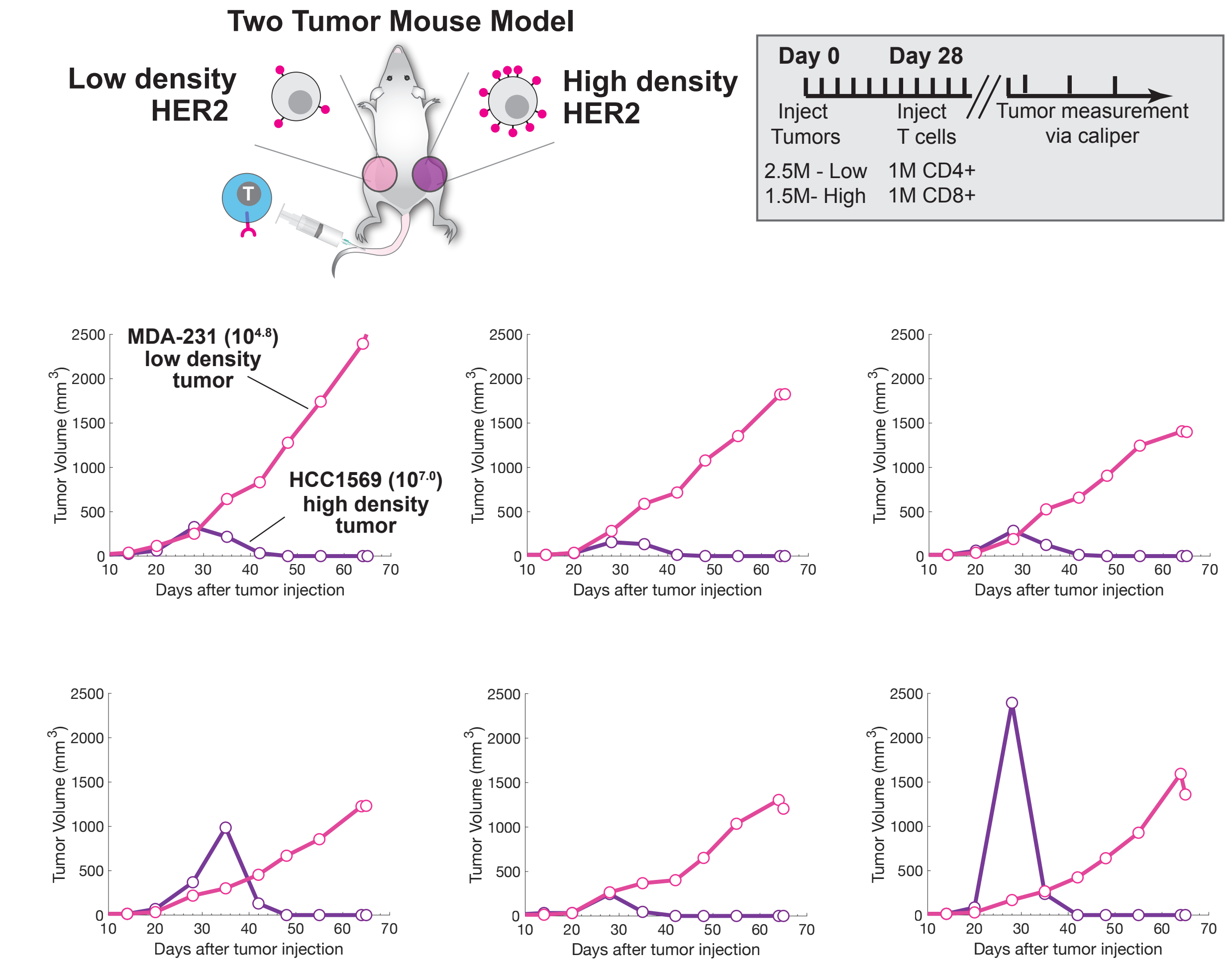


***B***

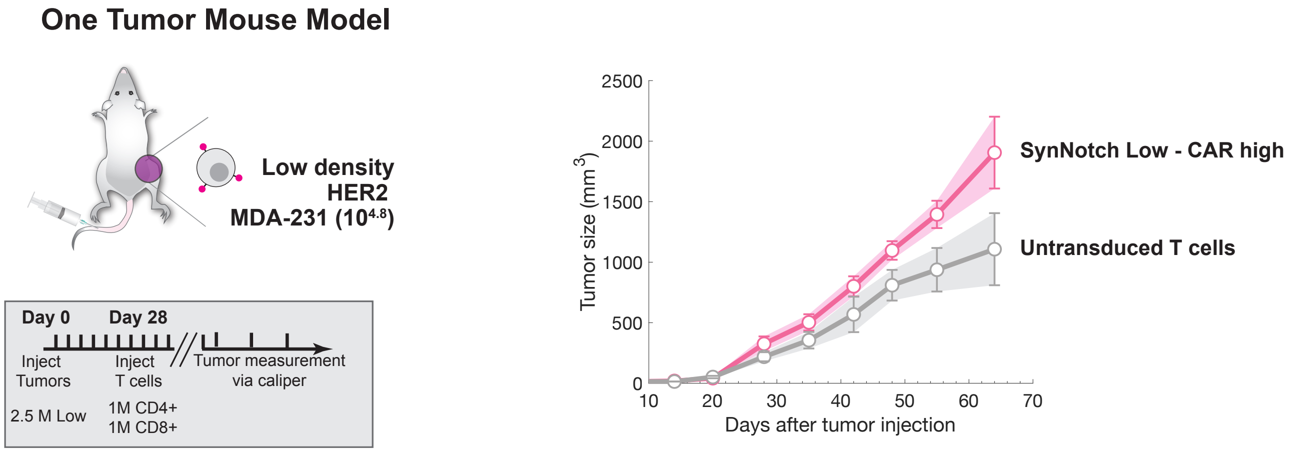

**Figure S8. Tumor volume measurements for individual mice treated with T cells expressing low affinity SynNotch to high affinity CAR circuit, related to Fig. 4D.**
**A.** Schematics of a two-tumor mouse model experiment. Low (MDA-231) and high (HCC1569) HER2 tumor cells were injected subcutaneously in the flanks of N.S.G. mice. Engineered primary human CD4+ and CD8+ T cells were injected i.v. 28 days after tumor injection. Tumor volume was monitored via caliper measurement over several days after tumor injection. Tumor volume data for individual mice treated with T cells expressing a low affinity SynNotch to high affinity CAR circuit. The dark purple lines correspond to the high (HCC1569) tumor whereas the light pink lines correspond to the low (MDA-231) tumor (See **Fig. 4D**). **B.** Schematics of a one-tumor mouse model experiment. Low (MDA-231) HER2 tumor cells were injected subcutaneously in the flank of N.S.G. mice. Engineered primary human CD4+ and CD8+ T cells were injected i.v. 28 days after tumor injection. The light pink line corresponds to the mean tumor volume for mice treated with T cells expressing a low affinity SynNotch to high affinity CAR circuit (n=4). The gray line shows the mean tumor volume after treatment with untransduced T cells (n=5).

**Fig. S9. T cells expressing a two-step circuit low-to-high SynNotch-CAR affinity recognition circuit yield ultrasensitive antigen density sensing against EGFR engineered cells.**

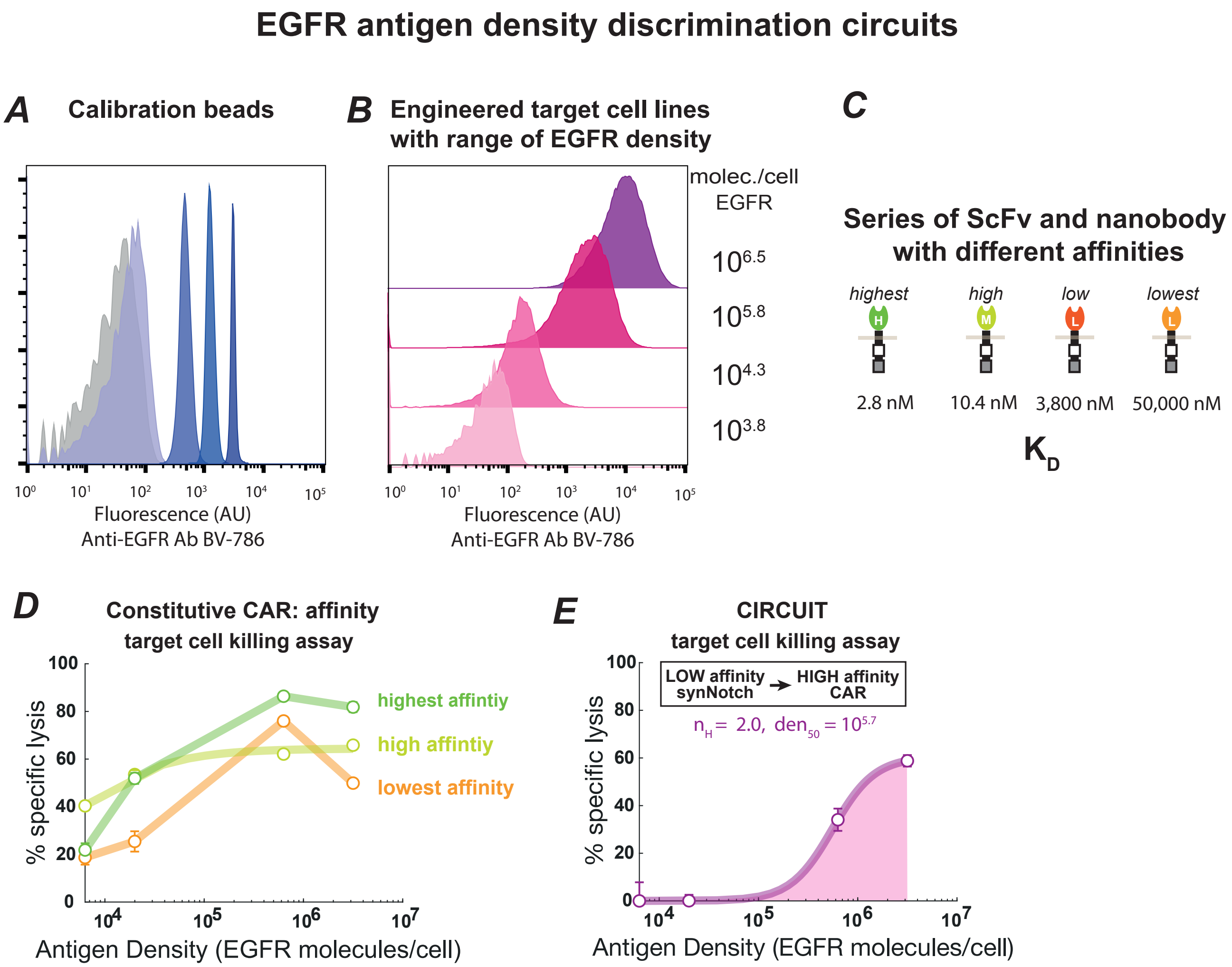


Targets (K562): 25,000
T cells (human primary CD8+):10,000
assay timepoint: 3 days

Targets (K562): 25,000
T cells (human primary CD8+): 30,000
assay timepoint: 3 days

**Figure S9. T cells expressing a two-step circuit low-to-high SynNotch-CAR affinity recognition circuit yield ultrasensitive antigen density sensing against EGFR engineered cells.**
**A.** Representative flow cytometry histograms showing the fluorescence intensity of Quantum Simply Cellular anti-Mouse IgG beads (Bang Laboratories 815) stained with anti-EGFR BV786 antibody, similar to **Fig. S1**. **B.** Representative flow cytometry histograms of engineered K562 EGFR cell lines stained with anti-EGFR BV786 antibody. The geometric mean of each population and the calibration curve built from data shown in the A was used to determine the average number of EGFR molecules per cell in each population. **C.** Series of scFv and nanobodies (*1, 2*)utilized to build two-step SynNotch to CAR circuits for EGFR density sensing. Their reported affinities are indicated. **E.** Target cell killing activity as a function of EGFR antigen density for T cells expressing CARs of indicated affinities. The E:T and assay time point are indicated. **F.** Target cell killing activity as a function of EGFR antigen density for T cells expressing a low affinity SynNotch to high affinity CAR circuit. The percentage of specific lysis was determined using flow cytometry by counting the number of target cells after 3 days relative to a co-culture in the presence of untransduced T cells. Data are shown as the mean and standard deviation from the mean (n=3). For the circuit transparent line shows a fit to a hill equation, the hill coefficient and antigen density for half maximal activity are indicated (see **Fig. S3B** for equation details).

**Movie S1.** T cells expressing a high affinity CAR (shown in blue) co-cultured with either low HER2 density cancer cells (PC3 shown in green) or with high HER2 density cancer cells (SKOV3 shown in red), imaged for 3 days. Still images shown in Fig. 3B.

**Movie S2.** T cells expressing a low affinity SynNotch to high affinity CAR circuit (shown in blue) co-cultured with either low HER2 density cancer cells (PC3 shown in green) or high HER2 density cancer cells (SKOV3 shown in red), imaged for 3 days. Still images shown in Fig. 3B.
